# Compositional Variability and Mutation Spectra of Monophyletic SARS-CoV-2 Clades

**DOI:** 10.1101/2020.08.26.267781

**Authors:** Xufei Teng, Qianpeng Li, Zhao Li, Yuansheng Zhang, Guangyi Niu, Jingfa Xiao, Jun Yu, Zhang Zhang, Shuhui Song

## Abstract

COVID-19 and its causative pathogen SARS-CoV-2 have rushed the world into a staggering pandemic in a few months and a global fight against both is still going on. Here, we describe an analysis procedure where genome composition and its variables are related, through the genetic code, to molecular mechanisms based on understanding of RNA replication and its feedback loop from mutation to viral proteome sequence fraternity including effective sites on replicase-transcriptase complex. Our analysis starts with primary sequence information and identity-based phylogeny based on 22,051 SARS-CoV-2 genome sequences and evaluation of sequence variation patterns as mutation spectrum and its 12 permutations among organized clades tailored to two key mechanisms: strand-biased and function-associated mutations. Our findings include: (1) The most dominant mutation is C-to-U permutation whose abundant second-codon-position counts alter amino acid composition toward higher molecular weight and lower hydrophobicity albeit assumed most slightly deleterious. (2) The second abundance group includes: three negative-strand mutations U-to-C, A-to-G, G-to-A and a positive-strand mutation G-to-U generated through an identical mechanism as C-to-U. (3) A clade-associated and biased mutation trend is found attributable to elevated level of the negative-sense strand synthesis. (4) Within-clade permutation variation is very informative for associating non-synonymous mutations and viral proteome changes. These findings demand a bioinformatics platform where emerging mutations are mapped on to mostly subtle but fast-adjusting viral proteomes and transcriptomes to provide biological and clinical information after logical convergence for effective pharmaceutical and diagnostic applications. Such thoughts and actions are in desperate need, especially in the middle of the *War against COVID-19*.

## Introduction

COVID-19, a novel pneumonia epidemic causing an outbreak first identified and reported in Dec 2019 from China [1] and subsequently spread to other countries swiftly, has been posing enormous professional, economic, and political challenges to global health services and hazardous control systems. As of 12 June 2020, there have been 7,410,510 confirmed cases and 418,294 deaths reported [2]. COVID-19 is of great contagious (even at incubation period) and has lower mortality to our current understanding [3-5]. The novel betacoronavirus identified through *de novo* sequencing from patients with COVID-19 is designated as “Severe Acute Respiratory Syndrome Coronavirus 2 (SARS-CoV-2)” by International Committee on Taxonomy of Viruses (ICTV) [1, 6, 7].

The recent threats from SARS-CoV-2, SARS-CoV, and MERS-CoV are different from those of earlier human coronaviruses (CoVs), including alphacoronaviruses, such as hsa-CoV-229E, hsa-CoV-NL63, hsa-CoV-OC43 and hsa-CoV-HKU1 [8-10], in at least two aspects. First, the recent groups of betacoronaviruses appears to come more frequently in the past two decades as compared to the early comers where new members may be discovered as technology become more efficient and accurate [11]. The current SARS-CoV-2 is also different from both SARS-CoV and MERS-CoV as its genome composition is most closely for “living with mammals and humans”, where a much lower G+C content has been evolved and is closer to two other human-adapted CoVs, hsa-CoV-229E and hsa-CoV-OC43, than its members of the recent group, although it shares higher sequence identities with the two new CoVs, 80.12% and 60.06%, respectively [11]. Second, it has been infecting far larger populations, as compared to the two recent outbreaks, with variable yet more complex symptoms [12]. The causative factors of such an unprecedented disease potency remain to be elucidated for the days and months to come [1, 3–7].

Genomes of coronaviruses mutate in a unique way where signatures of DNA pairing and repairing mechanisms are absent completely, and instead, they possess an error-prone synthesis of single-stranded full or partial genomic sequences by multi-component membrane-associated enzymatic structure known as the replicase-transcriptase complexes (RTCs) and double-membrane vesicles (DMVs) although they do have certain enzymatic activity resembling repair mechanisms of cellular organisms, such as proofreading [13], and other possible cellular mechanisms may also be involved, such as RNA editing as recently proposed [14,15]. Here we define a series of displays to understand compositional dynamics or variability that ultimately interconnects to proteomic variability including RTCs and DMVs (of course also other omics) through the organization of the genetic code [16–19]. We subsequently compare SARS-CoV-2 with other human CoVs for between-population variation analysis to point out that it is not a direct descendant of the previous human-infecting CoVs. We finally make efforts to decipher the SARS-CoV-2 clades for its variations and suggest that what we have seen now are not the natural picture of the pandemics and the missing-links are not among human populations but the wildlife close to human habitats in Southeast Asian territories, including islands and shorelines, not just limited to bats and pangolins. We also show how to examine clade-associated permutation variations and relate genetic variations to protein structures and phenotypic data. Nailing down a single animal of human origin of the virus will not be the goals of this genomics-based study but to provide information for smarter drug design, effective vaccine development, accurate diagnostics.

## Results

### Compositional dynamics and its parameters are essential and useful features for evaluating the evolutionary status and molecular mechanisms of SARS-CoV-2 towards pandemics

RNA genomics is very different from DNA genomics in several ways [11]. First, in the RNA genome, the A:U basepair is actually 2 Daltons heavier than that of the G:C basepair due to a larger molecular weight of uridine, whereas in the DNA genome the GC pair is a single Dalton heavier than that of the A:T basepair. In CoVs, the G+C content is actually in a trend of reducing so that the virus is in turn becoming heavier due to the increased U content [11]. Second, single-stranded RNA genomes are synthesized without stable double-stranded intermediates that allow mismatch repair albeit existence of short and extremely rare double-stranded RNA fragments involved in interference-based immunity [20, 21]. Third, the existence of Wobble basepairing for secondary structures is of essence for operational functions of all RNA molecules in addition to genetic information inheritance [22]. That said, we can now look at how the RNA genome of SARS-CoV-2 and related CoVs take advantage of these RNA-centric features.

As a positive-sense single-stranded RNA virus, SARS-CoV-2 has a genome length of ∼ 29903 nucleotides (nt) (GenBank: NC_045512.2). It encodes two large polypeptides, ORF1a and ORF1b, along with other 15 non-structure proteins (nsps; **Figure 1**A). In order to propagate and complete the life cycle, its positive-sense genome is first replicated to synthesize full-length negative-sense antigenomes and 10 shorter subgenomes (sgRNAs), executed by RTCs and DMVs, and the sgRNAs encode four structural proteins (S, spike; E, envelope; M, membrane; and N, nucleocapsid) and six accessory proteins (ORF3a, ORF6, ORF7a, ORF7b, ORF8, and ORF10) arranged among structural proteins depending on the current annotation (GenBank: NC_045512.2).

**Figure 1.**
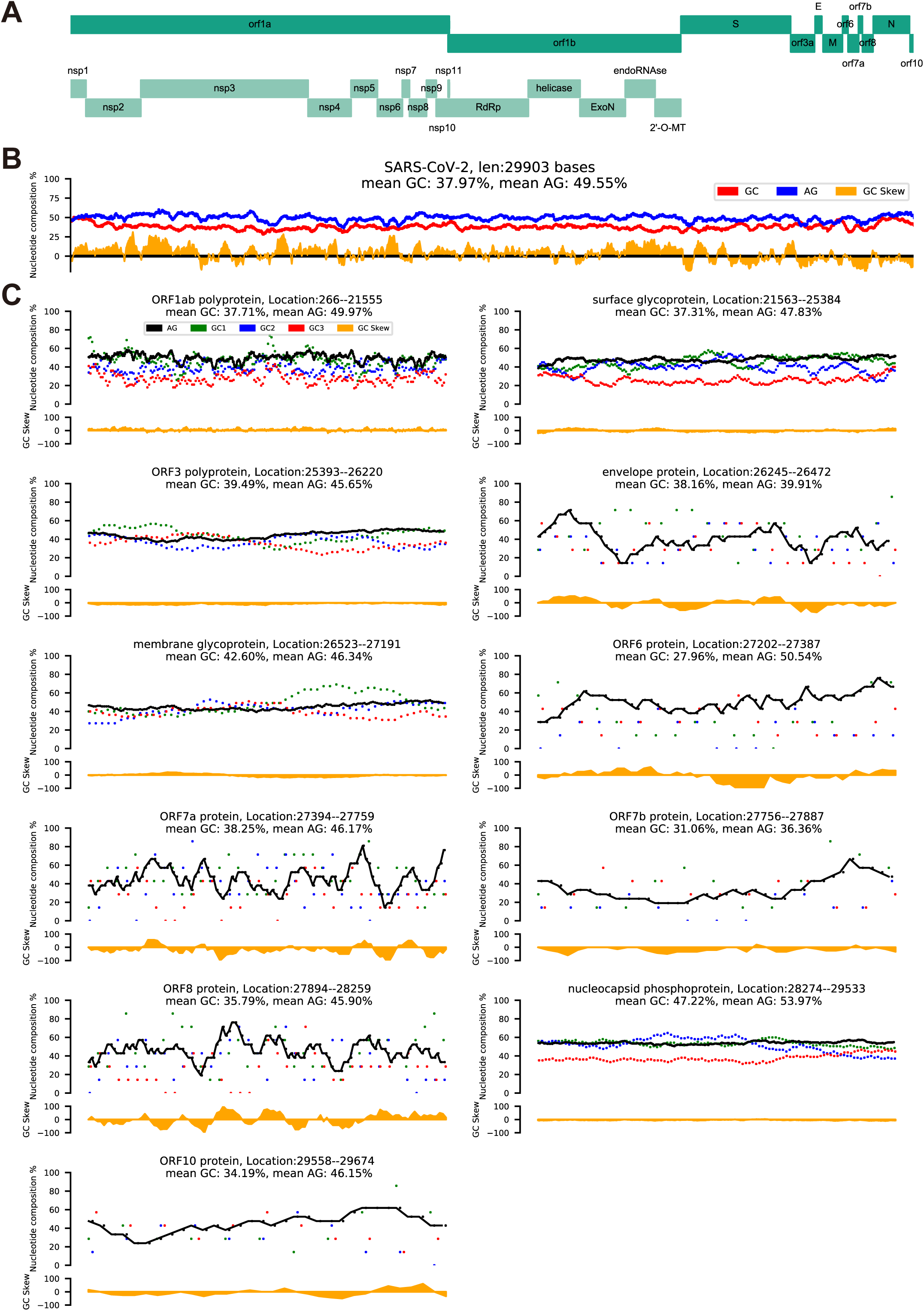
A display of genome compositional dynamics of SARS-CoV-2 and related CoVs. **A**. The complete genome sequence of SARS-CoV-2 (NC_045512.2), including both structural and non-structural components. **B**. We use a 300-nt sliding window with a 21-nt step to show dynamic changes of genome G+C, purine, GC1 to GC3, and GC skew (G-C/G+C) contents over the entire genomes. **C**. A similar procedure as described above is applied to individual ORFs and proteins. Note that the GC skews are not uniform over the genome length and the observation suggests possible recent recombination among closely related CoVs.

Traditionally, we use three basic plots to display composition dynamics based on primary genomic parameters over genome length: G+C contents at three codon positions (GC1, GC2 and GC3); purine content (A+G content); and GC skew (the content of (G-C)/(G+C)). Here, we use a 300-nt sliding window with a step size of 21 nt, as the majority of viral sequences are protein-coding, to illustrate the dynamics of the composition parameters, G+C and purine contents (Figure 1B). The G+C content of SARS-CoV-2 varies in a narrow but significant window of 18.00% (31.67%–49.67%) and the purine content in a slightly narrower window of 15.50% (41.67%–57.17%) in average over the entire genome length. The GC skew of the SARS-CoV-2 genome indicates the G+C ratio is relatively higher in structural proteins than ORF1ab and this imbalance is a signature of distinct mutational biases caused by viral replication machinery, known as RTCs. It is also variable as a frequent shift toward negative values are often seen in individual ORFs and defined proteins. Such minor anomalies suggest either recombination or selection events, which are species-or isolate-specific. The differences become obvious when SARS-CoV-2 is compared to the closely-related bat and pangolin CoVs (raf-betaCoV-RaTG13 and mja-betaCoV-P4L; Figure S1A and S1B), and the authentic within-species variation is exemplified when SARS-CoV and MERS-CoV are matched up with civets and camels in similar parameters, respectively (Figure S1C, S1D, S1E and S1F). The G+C content of different codon positions is also very informative, where GC3 is very characteristic of mutation pressure as it is obvious that all GC3 values of the viral proteins are biased toward lower G+C contents. GC3-associated mutations often reflect directional mutation patterns as observed strongly in certain lineages of plants and warm-blooded vertebrates as negative gradients from the transcription starts, and such trends are attributable to a special DNA repair mechanism, transcription-coupled DNA repair [23–25]. The notion here is to remind ourselves that transcription-centric mutations may be accounted for some of the mutation events in RNA viruses in their replication-transcription processes. Occasional twists from the trend often indicate selective pressures, such as in the case of S, M, and N proteins, and weaker GC3 or stronger GC1 or GC2 selections. Codon-associated G+C content trends are less informative for small ORFs, such as the case of ORF10. Most of the sequence signatures are indicative rather than proven functional relevance of proteins but very useful for providing clues of sequence anomaly.

For studying RNA viral genomes, in addition to previously-defined parameters, we need to introduce the concept of single nucleotide (A, U, G and C) contents at three codon positions (such as U1, U2 and U3 for uridines) (**Figure 2**) and to plot out compositional dynamics for positive-sense or the genome and negative-sense strands, which include both templates for synthesizing new positive-sense genomes or antigenome and subgenomes. All viral mRNAs are transcribed from an antigenome and other 9 subgenomes (Figure 2B). For compositional dynamics of RNA genomes, uridine is the star nucleotide, and A+U content becomes the most important. Just for the sake of convenience, we would like to keep the concept of G+C content since it has been known to be a useful variable for DNA compositional dynamics [26] and provide an approximation for less selected nucleotide position. For both DNA and RNA viruses, this variable G+C content remains similar to their hosts mostly, except those that are not well adapted to their hosts, such as SARS-CoV or MERS-CoV (Figure S2A; [11]).

**Figure 2.**
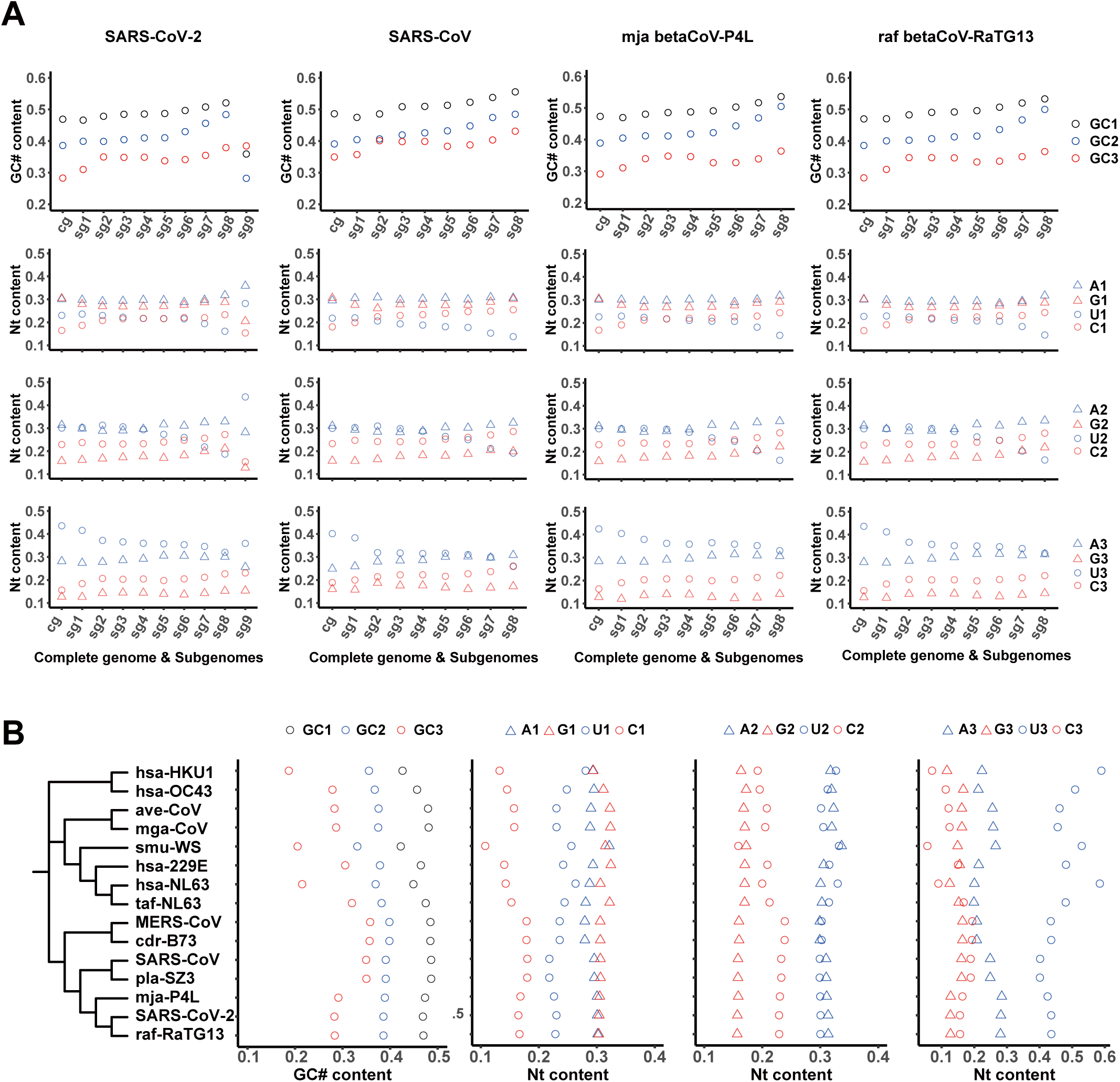
Nucleobase contents of genomes and subgenomes of SARS-CoV-2 and related CoVs. **A**. A schematic phylogenetic tree is used to cluster genome sequences and compositional variables (15 CoVs genome sequences, from top to bottom, are: hsa-betaCoV-HKU1, hsa-betaCoV-OC43, ave-gamaCoV, mga-gamaCoV, smu-alphaCoV-WS, hsa-alphaCoV-229E, hsa-alphaCoV-NL63, taf-alphaCoV-NL63, MERS-CoV, cdr-betaCoV-B73, SARS-CoV, pla-betaCoV-SZ3, mja-betaCoV-P4L, SARS-CoV-2, and raf-betaCoV-RaTG13). These compositional variables include GC contents at three codon positions (codon positions 1, 2, and 3 are denoted as GC1, GC2, and GC3) and single nucleotide contents at three codon positions (A1, A2, A3; U1, U2, U3; G1, G2, G3; and C1, C2, C3). Nucleotides are labeled in different shapes: purines, triangles; pyrimidines, open circles. A and U or G and C are colored blue or red, respectively. It becomes obvious that the two closely-related CoV genomes to SARS-CoV-2, the reported bat (raf-betaCoV-RaTG13) and the pangolin (mja-betaCoV-P4L) have very similar codon G+C content as well as base contents. The cp1 (codon position 1) base content appears most characteristic of balanced purine content of SARS-CoV-2 and its close relatives. The cp2 (codon position 2) base content of SARS-CoV-2 and all other CoVs has higher and relatively balanced A+U content. The older human CoVs have either lowest or higher G+C content and unbalanced purine contents. Note that G+C contents in three codon positions are labeled differently from those of single nucleotide contents in color codes. G+C content represents a single measure but single nucleotide contents demonstrate trends of all four nucleotides. **B**. The G+C and single nucleotide contents at different codon positions of complete genomes and subgenomes of SARS-CoV-2, SARS-CoV, mja-betaCoV-P4L, and raf-betaCoV-RaTG13 are displayed to illustrate the driving force for G+C content decrease toward 3’ end of the genome, which is rather a result of, in terms of mechanism, the increased U content and C-to-U permutation. The negative gradient of U is also obvious from the 5’ end to the 3’ end.

As shown in a phylogenetic tree constructed based on 15 representative coronaviruses (Figure 2A), the nucleotide content of SARS-CoV-2 is most similar to those of raf-betaCoV-RaTG13 and mja-betaCoV-P4L, which are considered to be distantly related but most closely related so-far-found host of SARS-CoV-2. Other known zoonotic and corresponding human counterpart CoVs are rather close to each other in their compositions. We have made a few interesting observations here. First, the single nucleotide content is more informative than G+C content, especially for genome analysis on RNA viruses. The former points out only how G+C content drifts toward richness or poorness but the latter narrows it down to single nucleotide effect. In our case, U stands out at cp3, which alters the overall nucleotide contents, and it drives the G+C content so low that even its partner A content has gone to the same extremity, so that the low G+C content is a result of both lowering U and A. If the organization principles are considered here, half of the codons are not sensitive to cp3 changes, and most of them are smaller amino acids (Figure S2B; [16–19]). Second, at the cp1, G and C contents are both pulled apart toward extremity but not A or U, while the two pyrimidines and two purines appear stretched to separate directions; these trends suggest strong selective pressure at the first codon position over the entire genome. It is indeed that cp1 codons shoulder the most mutation pressures since they fall into all 4 negative-sense strand permutations (known as R1-derived permutations, C-to-U, G-to-A, U-to-C and A-to-G). Third, the cp2 contents are most row-flipping changes referenced to the genetic code organization [18]. These alterations are very useful for alternating chemical characteristics between related amino acids, and in terms of flexibility, cp2 codons are less stringent than cp3 but more flexible than cp1. The balancing power becomes more obvious when ORFs or proteins are examined individually for their composition dynamics (Figure 1C). Finally, it is conclusive that the more similar the CoVs in composition dynamic parameters, the closer they are genetically and phylogenetically in principle. However, primary parameters, such as G+C and purine contents are necessary but may not be sufficient. For instance, there has been a CoV genome isolate from a wild vole captured in northeastern China, whose G+C and purine contents overlap with SARS-CoV-2 completely (Rodent coronavirus isolate RtMruf-CoV-2/JL2014; 0.38, 0.496; [27]) but its genome sequence is different (sharing 61.87% identity with SARS-CoV-2). Therefore, we have yet to find a within-population immediate animal host of SARS-CoV-2 albeit best similarity of composition dynamics seen among them.

Our subsequent study is focused on composition dynamics within CoV genomes. It is interesting to see uniformity among all codon position contents of all CoV genomes, increased G+C content from antigenomes to subgenomes. However, this trend is an illusion where the real trend is the lower G+C content of antigenomes but higher G+C contents of subgenomes due to stronger selection over structural proteins. This observation becomes clearer when all ORFs and proteins are scrutinized one by one (Figure 1C and Figures S1). SARS-CoV-2 has an exceptionally short subgenome 9 (sg9) which only contains ORF10, but we have no evidence that it is either functional or non-functional. These results collectively remind us that SARS-CoV-2 and its two most-closely-related CoVs, unlike in the case of many other known CoVs, have a unique genome composition and similar dynamics to the early-adapted human CoVs [11], and CoV-borne bats and other mammals may already coexist with ability to jump on to humans and domestic animals but only limited by environmental and geographic constraints.

### Mutation spectrum is composed of permutations that are distinct according to their strand specificity, order of synthesis, and ratio of positive-sense vs. negative-sense strands during propagation

We use 12 permutations to represent directional mutations and classify them according to strand-specific replication mechanisms (**Figure 3**) since they are readily related to codons [11] (Figure S2B). From a total of 5,054 point mutations, 1,416, 1,497, and 2,141 mutations fall on codon position 1, 2, and 3, respectively. The permutations are categorized into R1 (C-to-U, G-to-A, U-to-C and A-to-G), R2 (C-to-A, U-to-G, A-to-C and G-to-U), and R12 (C-to-G, U-to-A, A-to-U and G-to-C) derived according to their occurrence tailored to RTC-directed strand synthesis: R1 from the first negative-sense strand, R2 from the subsequent positive-sense strand, and R12 from R1 plus R2. The most abundant permutations are four R1 permutations and one R2 permutation, G-to-U (Figure 3A and 3B). What have we shown here is how sensitive are nucleotide content of cp3 to selective pressure, and most cp3 permutations disappear except the R2 G-to-U permutations at cp3, where all changes are transversions and more than half of all codons (all pro-diversity changes) are sensitive to them. Similar results are observed in our analysis on SARS-CoV and MERS-CoV (Figure S3A and S3B). There are slightly different patterns among SARS-CoV and MERS-CoV and their within-population mammals from the SARS-CoVs and close relatives, the higher U-to-C permutations. The predominate C-to-U represents a driving force of variation, and it manifests why both G+C and A+G contents of SARS-CoV-2 appear relatively lower against MERS-CoV and SARS-CoV and even more when compared to human CoVs, such as 229E and OC43 (Figure S2A).

**Figure 3.**
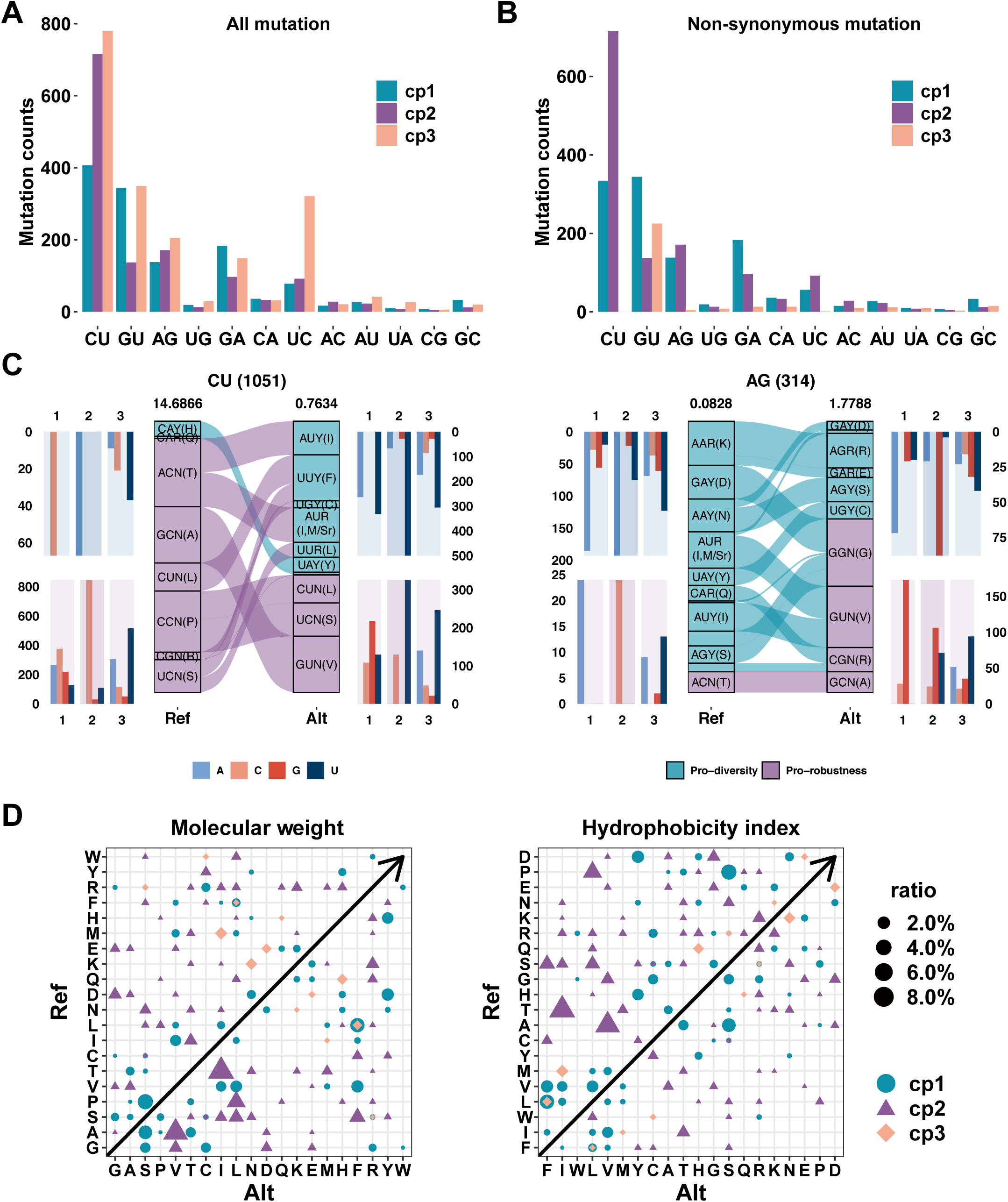
Mutation spectra of SARS-CoV-2 in a context of G+C contents and codon positions. **A**. The SARS-CoV-2 mutation spectrum is composed of 12 permutations and they are divided by codon positions among all mutations. C-to-U (CU), U-to-C (UC), A-to-G (AG), G-to-A (GA), and G-to-U (GU), are always dominant due to two principles; one is that the first four permutations occur when positive-sense genome is synthesized, and the other is that a G-by-A replacement is always preferred by RTCs so that G-to-U permutation as the most dominant occurring when the antigenome serves as a template. **B**. C-to-U permutations at cp3 diminish among non-synonymous mutations and this phenomenon indicates that most protein composition relevant variations are cp1 and cp2 variations. The remaining non-synonymous mutations in G-to-U (GU) permutation may be a result of biased strand synthesis. **C**. Displays of permutation-to-codon changes among non-synonymous mutations. The codon table is divided into two halves: the pro-diversity half (blue) whose cp3 is sensitive to transitional change and the pro-robust half (purple) whose cp3 position is insensitive to any change. Two examples, C-to-U (1051 in counts) and A-to-G (314 in counts) permutations are shown here. When a codon has a C-to-U change, the codon position varies, results of such changes relative to codon positions are summarized on both sides of the codon flow chart. Note that cp1 and cp2 changes appear more than those of cp3. The ratio between codons of pro-robust half and pro-diversity half is displayed on each bar. **D**. All permutations are plotted against the reference genome sequence to show how changes are related to amino acids. In the molecular weight index, most cp1 and cp2 changes are most obvious, showing an increasing trend. In the hydrophobicity index, most cp1 and cp2 changes increase toward less hydrophobicity.

Since most cp1 and cp2 related permutations are sensitive to selection, we have examined how individual permutations correlated to codon rearrangements in the two halves tables: pro-diversity and pro-robustness (Figure 3C and 3D) [16, 17]. Only two examples, C-to-U and A-to-G, are shown here and the rest are summarized in Figure S3C. Several observations are worthy of in-depth discussion. First, it is known that three amino acids and their codons are unique in balancing one of two purine-sensitive halves; they are Leu (leucine), Arg (arginine), and Ser (serine) [16–19]. The most abundant amino acid in protein coding sequences (known as codon usage) is Leu and it buffers C-to-U|U-to-C mutations at cp1. Arg and Ser are also abundant as they both are 6-fold degenerate codons; Arg appears buffering A-to-G|G-to-A at cp1 and Ser carries two: U-to-A|A-to-U at cp1 and G-to-C|C-to-G at cp2. Second, amino acid exchanges are permissive in physiochemical properties [23–25]. For instance, Ser has a very similar size to Ala (alanine) so that G+C content increase is buffered by the two amino acids as G-to-U|U-to-G permutations. Third, other examples are codon alterations among hydrophobic amino acids as they are mostly C-to-U changes at cp2 among those in the pro-robustness half. The overall effects are displayed together in Figure 3D. It is rather clear that changes toward lower G+C content and near the balanced purine content are both beneficial for CoVs, especially SARS-CoV-2, as these changes are pro-diversity, in favor of larger and more hydrophilic amino acids.

### Clade-associated biased mutation trend in SARS-CoV-2 revealed physiochemical features of replication machinery

Difficulties for analyzing CoV genomes are multifold. Since we have yet to identify the natural hosts and mammalian intermediate hosts, if there is any, this massive dataset has to be analyzed by stratifying the data into structured and non-structured clades; the former can be analyzed first and the rest await further ideas. The next is even more troublesome. Assuming that we have 5 or more genome sequences per CoV isolate and variations identified among them are still a miniscule fraction of the total virions produced in a patient body (medians and means of variations per CoV isolate among C01 to C09, see Table S3), since the viral load per patient sample, such as sputum [28, 29], is equivalent to a 5-person or more sampling of the entire human population on earth, 1 out of 10^9^. Even so, we have still been able to find shared variations among patient samples occasionally and even more lucky to have some clade structures, by and large due to the relatedness of the patients in the transmission network. Finally, we have to admit that many assumptions have to be made about these samples and their genome sequences above sequence and assembly errors for phylogeny and genetic studies. Nevertheless, we have constructed a somewhat stable phylogenetic tree-and-branch structure for further analysis (**Figure 4**A). It is composed of 8 monophyletic clades and 1 non-monophyletic clade based on both orders of sample collection date and highly-shared mutations. Among the clades, C02 shares two landmark mutations, C8782U in ORF1ab and U28144C in ORF8, and earlier date (2019/12/30). C04 shares three more mutations (C17747U, A17858G, and C18060U in ORF1ab) than what C02 have, and a late collection date (2020/02/20). Clades C03, C05, and C07 are also distinguishable by some major mutations, so are C06, C08, and C09; the latter clades are clustered together based on four shared and other clade-associated mutations. The leftover large number of isolates that lack all landmark mutations are grouped into C01, which have the earliest collection date on 2019/12/24. According to the literature and our discussion, we have further grouped the clades into three clusters, S (C02 and C04), G (C06, C08 and C09), and L (all the rest) since phylogeny shows clear divergence among them. We have several notions about this imperfect hierarchical structure. First, our within-and between-clade analysis of high major allele frequency (MAF) variations reveals that some clade-associated signature mutations are also shared among clades. For instance, C14805U in ORF1ab and A24034G in Cluster S have recurred in other clades of different clusters, which are excellent landmarks for subclade definition. Another notion is that higher MAF within-clade mutations (such as MAF>0.2) are mostly non-synonymous mutations, indicating selection at work (Figure S4). Our neighbor-joining tree based on distances from 9 clades suggests that SARS-CoV-2 appears originated from multiple zoonotic reservoirs instead of a single direct ancestor (Figure S4). In addition, our classification rationales are largely in agreement with published reports [30]; for example, Cluster S is in accordance with previously defined S type [31] and Cluster G is in line with GISAID [32] defined the G clade. Cluster L is similar to the V and L clades combined, of GISAID. A maximum likelihood (ML) based unrooted phylogenetic tree is shown in Figure 4B.

**Figure 4.**
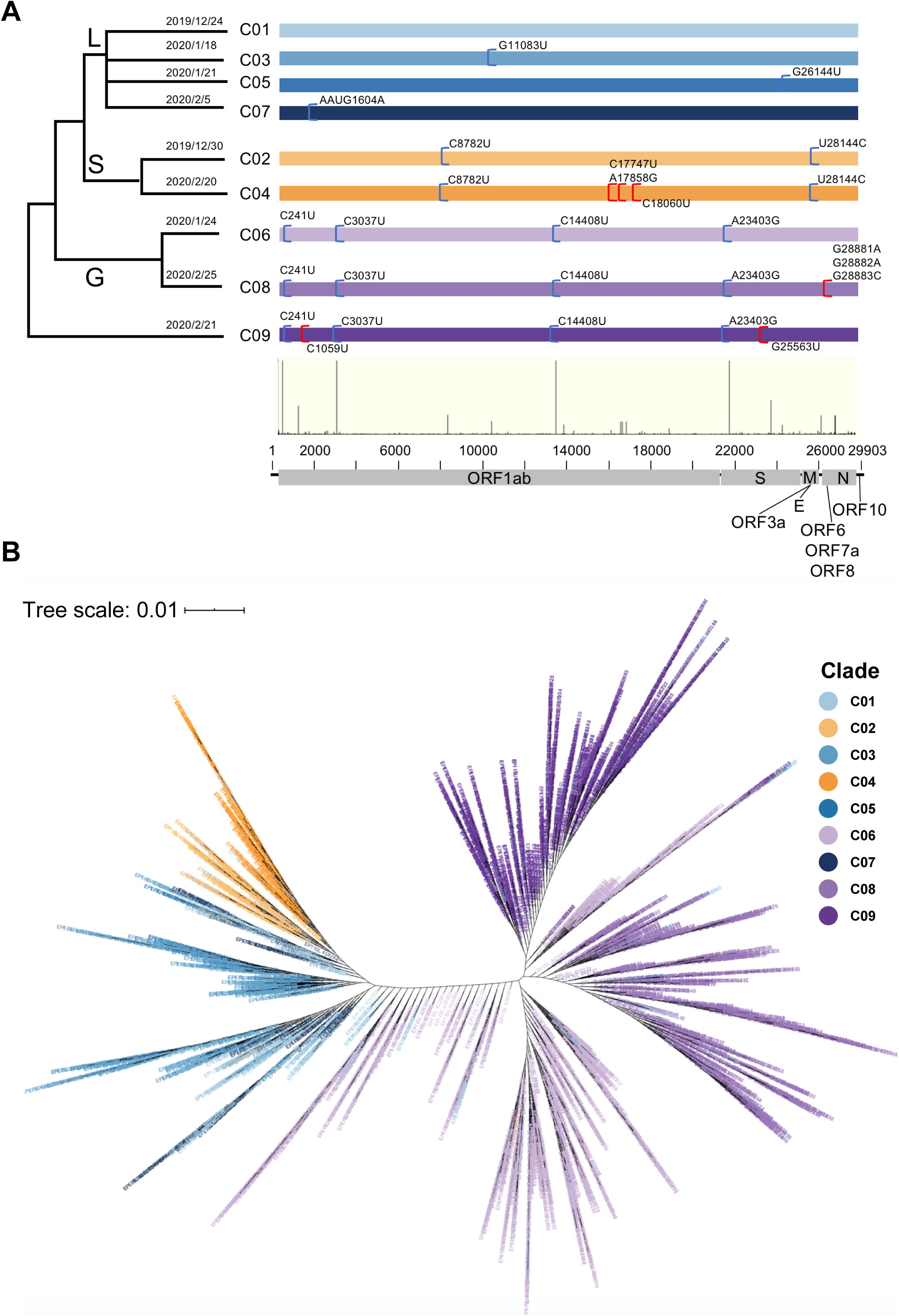
Sequence-variation-based phylogenies of SARS-CoV-2. **A**. CoV genomes are divided into clades and clade clusters based on high-frequency mutations among the genome sequences. The shared variations are excellent indicators for shared ancestors and those between clusters (blue half parentheses) and within clusters (red half parentheses) are labeled with positions and nucleotide variations that are all referenced to the SARS-CoV-2 genome (NC_045512.2), its positions, and relative frequencies (thin vertical bars). The dates when each clade started are also indicated. **B**. The current collection shows 9 clades (C01 to C09) in three clusters (S, L and G). An unrooted phylogenic tree of the clades and clusters (color-coded), the tree scale is 0.01.

To look for clade-associated compositional and functional features, we have first built a consensus sequence for each clade and subsequently calculated frequencies for each within-clade permutation (Table S2; **Figure 5**A and 5B). A key assumption behind this is that certain functional mutations may have clade-specific effects on mutation spectrum, to close a loop where sequence mutations through genetic coding principles alter the viral proteome function. Our observations are of importance in establishing logics about compositional dynamics between nucleic acids and proteins. First, permutations among clades are indeed variable according to their proportions calculated from genome variants, and aside from 5 high-proportion permutations, 4 R1 and 1 R2 permutations, two other R2 and one R12 permutations appear also joining in, which are U-to-G and A-to-C, as well as A-to-U, respectively. Second, the variable permutations, where some may represent effect of mutation pressure and others may exaggerate selection pressure, are unique to clades and clade clusters. For instance, clade cluster S has the lowest G-to-U fraction as compared to those of L and G; in addition, among the S clades, C04 has the lowest value of G-to-U. Similarly, C03, C05, C06, C08, and C09 have relatively higher G-to-U permutations. Third, based on the disparity of permutations or simply mutation spectra, we have taken a rather radical step to assume RTC statuses in favor of either *tight* or *loose* statuses for binding to purines and pyrimidines (see Figure S5). Since purines are larger than pyrimidines in size, the purine-or R-tight must be different from pyrimidine-or Y-tight. The results are strikingly predictable in that the R-tight status suggests a tighter binding pocket where a descending trend for tight permutations (C-to-U, G-to-U, and U-to-A) reverses into the opposite trend for Y-tight permutations. It indicates that the RTC structure and conformation variables may be definable in principle. At this point, we do not have discrete definitions for these so-called tight statuses but the less trendy R-loose and Y-loose statuses also support a similar idea.

**Figure 5.**
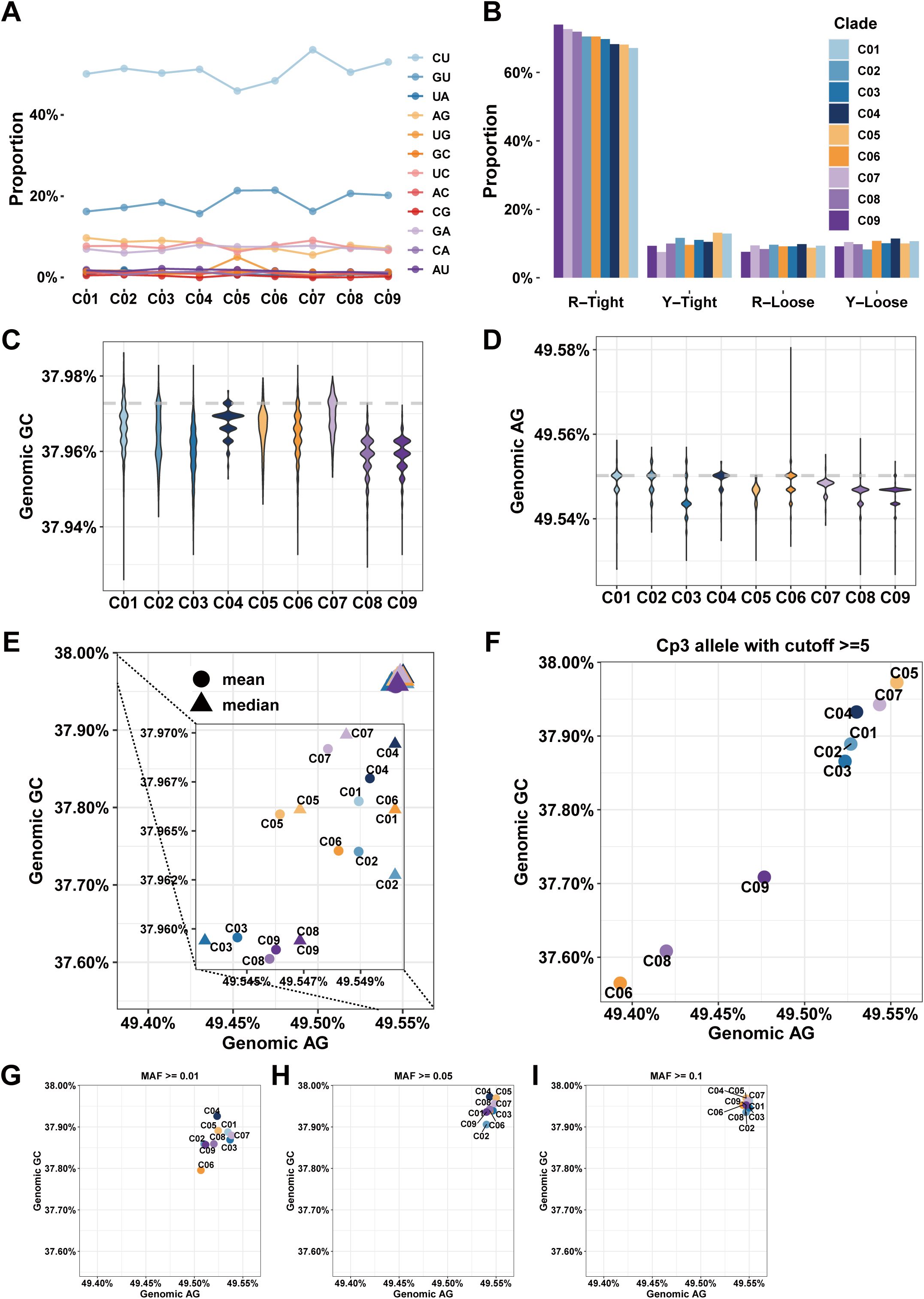
Mutation spectrum and composition dynamics among 9 SARS-CoV-2 clades. **A**. Plots showing permutation variation of each clade. Aside from the 5 dominant permutations, A-to-C (R2 permutation), U-to-G (R2 permutation) and A-to-U (R12 permutation) changes appear also significant; such an increase in proportion of R2 and R12 permutations often indicates copy number (synthesis) bias between the two strands. **B**. When permutations are grouped based on structure-conformation model (Figure S5) into tight and loose groups (a four-parameter model), their trends of changes become obvious. The R-tight discourages A-by-G replacement but encourages C-by-U replacement when the genome is replicated. The loose statuses, regardless R-loose or Y-loose, place no pressure on permutation variability. **C**. Violin plots showing the G+C content among clades. **D**. Violin plots showing the purine content among the clades. C08 and C09 have been drifting both contents toward lower ends. C03 has also been drifting in a greater extent of its purine content and a lesser extent of its G+C content, comparatively. **E**. The mean (solid circles) and median (solid triangles) of G+C and purine contents among clades. The same three more expressive clades, as seen in (**C**) and (**D**), are indeed obvious (inset). **F**. The compositional dynamics of cp3 nucleotides that are less selected and with a stringent cutoff value (> or =5). **G**. Composition distributions based on major alleles, at frequencies equal or greater than 0.01 to emphasize the effect from selection. **H**. Composition distributions based on major alleles at frequencies equal or greater than 0.05. **I**. Composition distributions based on major alleles at frequencies equal or greater than 0.1.

We have further examined the compositional subtleties among the clades and clusters with a focus on G+C and purine content variability as both contents appear drifting toward optima in SARS-CoV-2 and its relatives (Figure 5C and 5D). Different clades exhibit distinct compositional features and such dynamics are very indicative for the existence of feedback loops connecting RNA variables to protein variables. Two directions have to be advised for understanding these features albeit in absence of between-clade statistics. The first direction is driven by strong mutations, perhaps coupled to tight-loose switches in the catalytic pocket of RdRPs in RTCs. It is clear that except C01, the G or C06-C08-C09 cluster has the lowest G+C (0.37929, based on a C08 CoV sampled in Australia) and the lowest purine contents (0.49527, based on a C08 CoV collected in Bangladesh and a C09 CoV collected in England). Both lower G+C and purine contents are indicative of mutation pressure and signal this fast-evolving cluster of CoVs. Since this cluster has the largest collection of CoVs, it is also not surprising to see a more complex median diversification within clades (Figure 5C and 5D). The second direction is the drive from selection or both selection and mutation in balance or imbalance, as well as in modes of fine-tuning or quick-escaping. Some results from our analyses are shown here for briefing purposes (Figure 5E to 5G). For instance, G+C and purine contents at cp3 are informative for mutation drives and other measures are less clear cut (Figure 5F), given the evidence that even MAFs among clades are not stably distributed among clades as lower MAF variations are rather sporadic and hard to analyze even binned into groups (data not shown).

Based on our clade and clade cluster analysis, it is tempting for us to speculate that there are plenty of rooms for further investigations into mutation spectra among large clades and even smaller clades or closely related individual CoV genomes for several reasons. First, all high-frequency MAFs should be identified and classified and these variations are candidates for highly selected mutations. Second, all within-clade minor but not rare alleles (less than 1/10,000), such as those of MAFs in a range of 0.01% to 10% should also be identified; they provide basis of within-clade sequence analysis. Third, all non-structured CoV genomes must be also classified based on shared variations, as they are not only valuable for within-clade but also for clade-cluster analyses since there is a large background of genome variations not yet brought into the databases.

### Within-clade variations and their implications for future SARS-CoV surveillance

Within-clade compositional dynamics can also be very informative, especially for covering and predicting future functional changes, such as identifying mutated and diversified forms of CoVs for drug and vaccine designs. It is also of essence for nucleic acid-based diagnostics, such as clade-specific identifications. We are in a process of developing an interactive database and mutation-function predicting algorithms based on our results to interpret novel sequence variations in real time. Within-population variations are identified based on clade consensus sequence after alignment and extracted from datasets that have hundreds and thousands of genome sequences. The analysis of within-population variations relies on structured phylogeny and proportion change of permutations. The changes, based on functional relevance, can be classified into either copy number-related or RTC-specificity related, or sometimes both.

We have taken two steps to extract information in order to distinguish the underlining mechanisms (**Figure 6**). In the first approach, we identify key mutations based on MAF of mutations with a consideration of relatively even distribution among subclades and name the subclades in a sequential order based on the absence of a subset (Figure 6A). In the second step, we plot out permutations to track changes among subclades (Figure 6B). For instance, clade C02 can be divided into 8 subclades and its variable permutation fractions are clearly recognizable. An immediate discovery is the trends of descending C-to-U, ascending A-to-G, and wavy G-to-U that initially goes up with A-to-G but rides down with C-to-U afterward.

**Figure 6.**
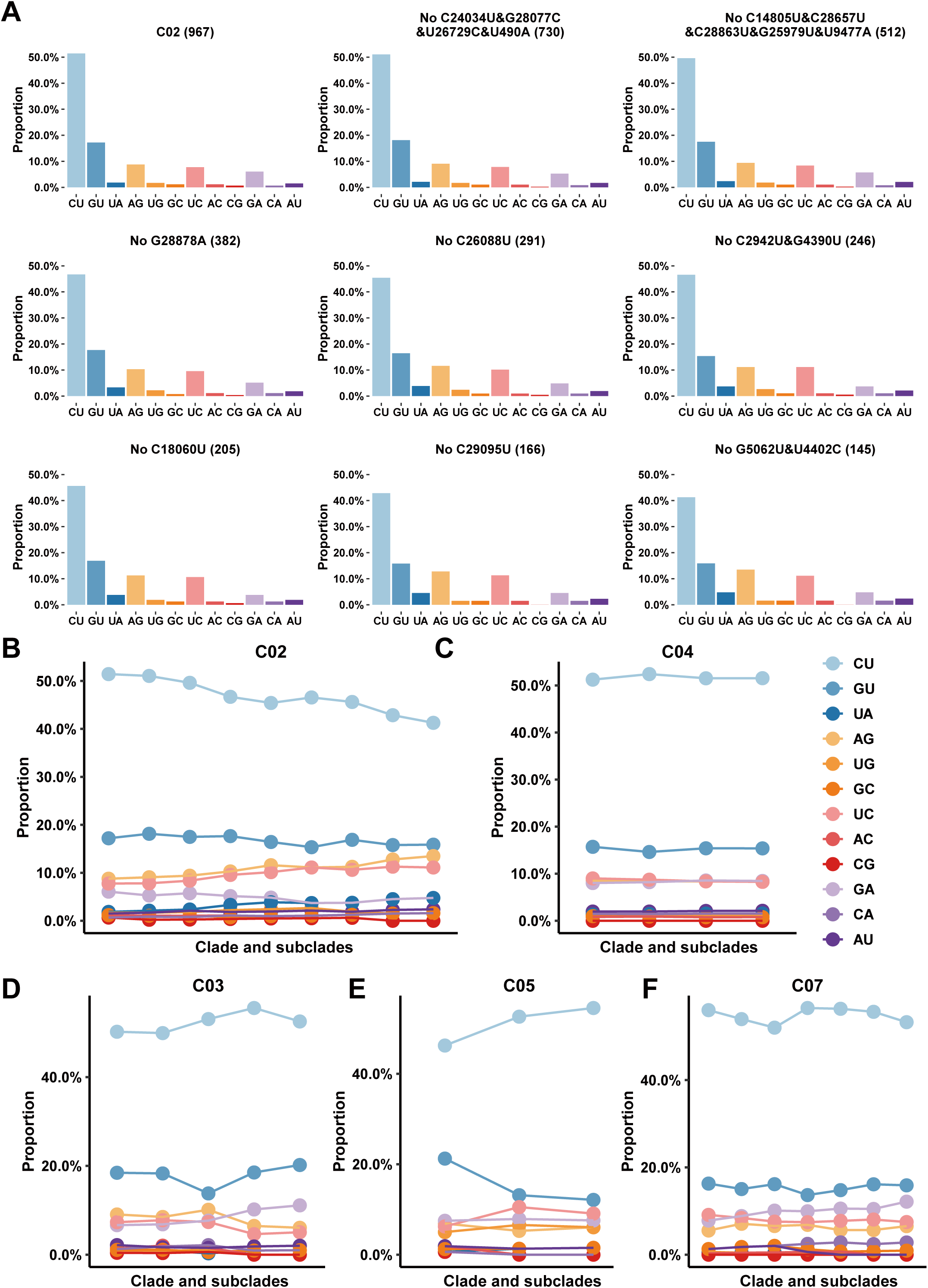
Within-clade permutation variations are excellent indicators of functional mutations. **A**. An example of permutation shifting of clade C02 and among its subclades. The number of SARS-CoV-2 genomes is indicated in the parentheses. The clear trends are two-fold. First, decreased C-to-U permutation is coupled with increased A-to-G and decreased G-to-U permutations. Second, A-to-U permutation is also increased as expected based on the model shown in Figure S5. These trends pf permutation changes suggest irrelevant to the ratio of strand-biased synthesis (positive sense vs negative sense) but possible structural and/or conformational variation in the RTCs. (**B)** – (**F)** show within-clade permutation changes of C02, C04, C03, C05 and C07. In each display, the first column of the x-axis shows the proportion of permutations calculated for each clade. Two opposite trends of permutation variations are seen between C03 and C04, and C07 has a rather wavy pattern.

Taking the two smaller clades, such as C03 and C05, as examples (Figure 6D and 6E), we first find that their trends of permutation variables show opposite directions, where the increasing C-to-U accompanies with the decreasing G-to-U. A closer examination reveals that the increasing C-to-U in C03 is also accompanied by descending U-to-C. The only permutation showing an increasing trend in C03 is G-to-A. The take-home message from these trends is that RNA synthesis of this subclade is biased toward producing more negative-sense strands or its mutation spectrum exhibits increasing mutations generated during the negative-sense strand synthesis. Such analysis can be carried out continuously when more CoV genome data become available as other within-clade variations are not as informative as C03 vs. C05 (Figure S6).

Several precautions are worth noting in such analysis. The most noticeable weakness is the fact that we assume function-related mutations are discovered in our dataset. As we have proposed an analogy before, chances are slim, dozens out of millions or even billions. Furthermore, even if we see drastic changes in permutations and mutation spectra, the mutations we identified still need validation empirically and based on different data types or sources albeit rare and precious. Finally, most frequently encountered situations are those that multiple mutations exhibit cofounding effects for a phenotypically identified functional or structural feature, and undoubtedly, more and deep-sequencing data are still invaluable and irreplaceable.

## Conclusion

This COVID-19 pandemic provides once-in-a-lifetime opportunity for the fields of biomedicine and other life sciences to work together on it as many facets as possible albeit exchanging with lives and other massive losses. If lessons told, we had learned things in serious ways in the last two CoV epidemics and we did prepare ourselves with vaccines and medication since, we would not have suffered this much this time. If one assumes that the last two outbreaks of SARS-CoV and MERS-CoV came surely by chances, this time SARS-CoV-2 is here for real, and a worst-case scenario is that it may stay with us forever or until effective vaccination is developed. Nevertheless, it certainly will stay with us for quite a while for many reasons [11]. First, at least it and other within-population versions of coronaviruses will definitely come again because we have not been able to trace its origin and ways its transmission from the very beginning, neither the Wuhan outbreak nor the recent Beijing outbreak in China even guided with very strict quarantine roles and prompt action plans. Next, this particular virus, SARS-CoV-2, has evolved to a composition status where some of its natural yet genetically distant hosts or possible intermediate mammalian hosts have acquired similar status [33, 34]. Furthermore, we do not yet have enough data to really map out the phylogenetic position that allows us to pinpoint its natural origin and human transmission routes.

The number one needs for us is data, genomic and clinical data, which should be as complete as possible and with characteristics including high-quality and high-coverage at single-molecule resolution. We currently have been acquiring genomic data and the specialized databases have collections over ten thousand non-redundant sequence variations, but still not enough to address more than a few possible functional changes of some key protein components [35–38], let alone understanding mutation-centric cellular mechanisms. Based on median and mean estimates, we have on average a mutation accumulation rate of half a dozen per patient. Although there have been data reported from single-molecule sequencing platform but they are low in coverage [39].

Our final notion is to emphasize the importance of analysis strategies and supporting platforms. Since questions always overwhelm what we can possibly address [40], prioritizing tasks are of essence together with choices of strategies. The first platform to be established concerns mutation-to-function interpretation, where we have present one in this report. Another to be considered is mathematic modeling, such as cellular and disease transmissions [41–46] and viral mutation-selection paradigm, for testing and evaluating different parameters and prioritizing what kind of data to be acquired with high priorities. In addition, cellular and molecular data, including different omics studies [47], all need to be incorporated into a COVID-19 knowledgebase, where information from multi-disciplinary studies are managed, organized, and mined.

## Materials and Methods

### SARS-CoV-2 and other related coronaviruses sequences

We used the public-available SARS-CoV-2 data collected worldwide among the major databases, including CNCB/NGDC [48], CNGBdb [49], GISAID [32], GenBank [50] and NMDC [51] on June 12^th^, 2020. To ensure authenticity and reliability, our datasets must meet the following criteria: (1) The genome sequence is labeled as complete that covers all coding regions of the reference genome (GenBank accession NC_045512.2). (2) It has no more than 15 uncertain bases that often substituted as “N”s. (3) It has no more than 50 degenerate bases that often labeled as discrete nucleotides. These high-quality genomes were aligned to the reference using MUSCLE (version 3.8.31) with default parameter settings [52]. Further analyses of SARS-CoV-2 and related CoV genomes are referenced to genome annotation of the same reference genome (NC_045512.2) and other information provided by the RefSeq database at NCBI.

Other closely related CoV genome sequences used include hsa-betaCoV-HKU1, hsa-betaCoV-OC43, ave-gamaCoV, mga-gamaCoV, smu-alphaCoV-WS, hsa-alphaCoV-229E, hsa-alphaCoV-NL63, taf-alphaCoV-NL63, MERS-CoV (from human and camel hosts), cdr-betaCoV-B73, SARS-CoV (from human and civet hosts), pla-betaCoV-SZ3 and raf-betaCoV-RaTG13 are retrieved from NCBI and mja-betaCoV-P4L are retrieved from NGDC. A full listing of our sequences dataset including virus genre, strain name, accession number and sources is provided in Table S3.

### Calculation of genomic composition parameters

We display several genomic composition dynamics and its parameters (G+C content, A+G content and GC skew) using different sliding windows. The first 300 nt are grouped as an initial window, and subsequent windows are uniformly shifted in a 21-nt step. Within these displays, the G+C contents referenced to the three codon positions of each open reading frame or ORF are measured by adjusting the sliding window according to the ORF lengths within viral genomes. As for ORFs longer than 2000 nt, a relatively large window size (300 nt) is adopted, and the step size is calculated via a custom formula 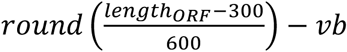, 0 ≤ *vb* ≤ 2 where *length*_ABC_ denotes the length of ORF and *vb* varied from zero to two bases to make sure the window size is divisible by 3; for ORFs with a medium size (longer than 500 bases and shorter than 2000 nt), the window size is defined as 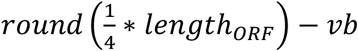, 0 ≤ *vb* ≤ 2, while the step size is simply defined as 3 nt; as for those small ORFs (shorter than 500 nt) such as structural proteins, a constant 21-nt window size and 3-nt step size is used for calculating genomic composition frequency.

The criteria for choosing the representative CoV genome sequences for constructing a representative phylogenetic tree (Figure 2) are multi-fold. First, we include all 7 human-infecting coronaviruses for the analysis, which are SARS-CoV, SARS-CoV-2, MERS-CoV, hsa-alphaCoV-229E, hsa-betaCoV-OC43, hsa-betaCoV-HKU1, and hsa-alphaCoV-NL63 (a prefix hsa-standing for *Homo sapiens* was used to label the unfamiliar human-infecting CoVs). Second, we categorized all human-infecting, for simplicity, into 4 lineages: SARS-CoV-2, SARS-CoV, MERS-CoV, and the older human CoV lineages. Therefore, their related CoVs in the literature were also selected for the analysis, including a single closely-related CoVs for each lineage (based on sequence identity): SARS-CoV-related (pla-betaCoV-SZ3), MERS-CoV-related (cdr-betaCoV-B73), SARS-CoV-2 related (raf-betaCoV-RaTG13), and NL63-related (taf-alphaCoV-NL63; both species and CoV genera were labelled for clarity). Third, we also added more informative CoV genome sequences to enrich lineage-associated information, which are a pangolin coronavirus genome (mja-betaCoV-P4L) reported to be closed to SARS-CoV-2 and 3 non-beta-coronaviruses that infect animals (e.g., ave-gamaCoV from gamma-coronavirus genus and smu-alphaCoV-WS from alpha-coronavirus genus). Fourth, we only used complete protein-coding sequences from the CoVs to construct the phylogenetic tree and to calculate genome parameter contents. The sequences were aligned by using MUSCLE and the UPGMA tree was constructed by using MEGA-X [53]. The G+C content and single nucleotide content of each virus genome at three codon positions was also calculated. Subgenomes of SARS-CoV was obtained from Marra et al [54], and we annotated the subgenomes of SARS-CoV-2, mja-betaCoV-P4L, and raf-betaCoV-RaTG13 based on the annotation of NCBI (GenBank accession NC_045512.2). In addition, G+C and single nucleotide contents of the complete genome and its subgenomes of these four viruses at three codon positions were displayed to serve as sequence composition references.

### Variation calling and categorization

All sequence variations are identified and categorized based on comparisons between the query and the reference genomes, and files were generated by using an in-hoc Perl scripts based on alignment results. The tailored annotation (gene, location and consequence on the protein sequence) of each variant is determined with VEP (version 99.0) [55]. Since a large number of gaps and low-quality sequences at the 3’ and 5’ ends, variations (substitutions, insertions and deletions or indels) occurring 50 nt each at 5’-and 3’-ends of the genome are not considered. Since the higher quartile of variations per genome among SARS-CoV-2 populations is 9 (based on the 22,051 sequences we analyzed in this study), we filtered out the problematic sites that exceed 50 variations as compared to the reference genome. CoV genome sequences have at least one mutation are used in this study. A full listing of variations among coding regions identified in this study is provided in Table S4.

All continuously updated mutation files of the SARS-CoV-2 populations in variant call format (version 4.2) are deposited at the variation page of the 2019nCoVR database contributed by CNCB/NGDC (https://bigd.big.ac.cn/ncov/variation/).

### Mutation spectrum analysis

A mutation spectrum for within-population variations is composed of two lines of information; one concerns mutations that are referenced to a population consensus built based on the entire collection, and the other contains frequencies of all mutations and their directional changes, i.e., permutations. To reduce pitfalls of sequencing errors, we only selected mutations that occur more than twice in the whole collection of SARS-CoV-2 populations (clades or clade clusters that are often defined based on phylogenetic analysis). In theory, there are 16 possible permutations but 4 of them are unrecognizable so that 12 permutations (C-to-U, A-to-G, U-to-C, G-to-A, G-to-U, U-to-G, A-to-C, C-to-A, U-to-A, G-to-C, C-to-G and A-to-U) are there as an informative set. When the number of CoV genomes collected are limited, such as SARS-CoVs and MERS-CoVs, entire data sets are pooled together without clades. In our analyses on SARS-CoVs and MERS-CoVs (Figure S3A), we aligned sequences from these two lineages to their reference genomes (SARS-CoV: NC_004718.3; MERS-CoV: NC_019843.3) to call variations. When aligned on overlapping sequences, due to large deletions and additional ORFs, are encountered, we always choose the largest or only one of the ORFs to represent the segment, respectively. For example, in SARS-CoV lineage, if a mutation falls into the overlapping region of ORF9a (encoding the N protein) and ORF9b, we have only used the ORF9a annotations to avoid redundancy.

### Phylogeny constructing

Given the scale of SARS-CoV-2 sequence collections, we focused on genomes with unique information contributing to phylogenetic analysis. First, mutations (including single-nucleotide substitution and indels) at frequencies equal or greater than 10 in between-clade or –population calls were selected. FastTree (version 2.1.11) [56] is used to construct maximum likelihood phylogeny based on 5,121 genomes that have met our criteria, and iTol [57], an interactive web server was employed for setting an unrooted format and annotating samples.

For Figure S4, the neighbor-joining method is used for constructing phylogeny from the Euclidean distance of the mutation frequency matrix of clades, and the tree was generated and visualized by R package phangorn [58] and ggtree [59].

### Estimation of G+C and purine contents of genome sequences

G+C and purine (or A+G) contents of CoVs in general vary in a narrow range, and therefore, subtleties among the content changes have to be scrutinized with low-quality sequences excluded. A more sensitive approach is used in this study where two points are assumed; all genomes are full-length and variant alleles in coding sequences are the varied composition. The absolute frequencies of A+G and G+C content are defined as:

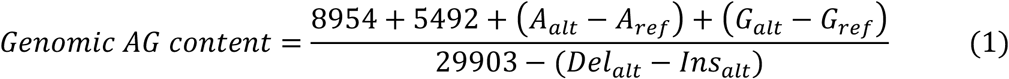

and

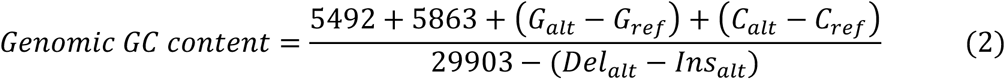

where 8,954, 5,492, 5,863, and 29,903 are the frequencies of A, G, C and total length of the SARS-CoV-2 reference, respectively. For any sequence compared with the reference, the *Del*_*alt*_ and *Ins*_*alt*_ measures the deleted and inserted nucleotides of this sequence, respectively, and that is why (*Del*_*alt*_ – *Ins*_*alt*_) means the variation of sequence length. For all the variant sites in this sequence, (*A*_*alt*_ – *A*_*ref*_) + (*G*_*alt*_ – *G*_*ref*_) in Equation (1) measures the number of A and G variations in compared sequence, *A*_*alt*_ and *G*_*alt*_ denote the number of nucleotides mutated to A or G while *A*_*ref*_ and *G*_*ref*_ represent the number of nucleotides mutated from A or G. Similarly, S*G*_*alt*_ – *G*_*ref*_) + (*C*_*alt*_ – *C*_*ref*_) in Equation (2) represents the varied number of G and C among compared sequences.

### Clade subgrouping

To detect trend followers and disrupters in mutation spectra, a pipeline was developed to select such genomes and mutations within clades iteratively. The first step includes locating high-frequency mutations (major alleles, MA) in a clade and extracting all genomes without this MA mutation to form a subset of the clade. The second step is, within the new subclade, to iterate the process until such mutations are thoroughly identified and no more mutations exceed a manually set threshold of MAF. Since the number of unique variations among clades have been varying significantly over time, the thresholds are 0.05 in C01, C04, C06, C08 and C09; 0.1 in C02, C03, C05 and C07. The proportion of permutations in each subclade and the located gene and mutation type (synonymous or non-synonymous) of mutations are provided in Table S5.

## Authors’ contributions

JY designed, supervised, and coordinated the study. SS, ZZ and JX participated in the design of the study. XT, QL, ZL, YZ, GN performed the data analysis. JY, SS, XT, QL, ZL, YZ designed and drew the figures. JY and XT drafted the manuscript. JY, ZZ, SS, QL and XT revised the manuscript. All authors read and approved the final manuscript.

## Acknowledgements

This work was supported by grants from The Strategic Priority Research Program of the Chinese Academy of Sciences [XDA19090116 to S.S., XDA19050302 to Z.Z.], National Key R&D Program of China [2020YFC0848900, 2017YFC0907502], 13th Five-year Informatization Plan of Chinese Academy of Sciences [XXH13505-05], K. C. Wong Education Foundation to Z.Z., and International Partnership Program of the Chinese Academy of Sciences [153F11KYSB20160008]. The Youth Innovation Promotion Association of Chinese Academy of Science [2017141 to S.S.]; Funding for open access charge: The Youth Innovation Promotion Association of Chinese Academy of Science. We thank our colleagues and students for their hard working on the 2019nCoVR (https://bigd.big.ac.cn/ncov). We thank Dr. Lina Ma, Lili Hao, and Meng Zhang for useful suggestions and discussion of this manuscript.

## Supplementary material

**Supplementary Figure S1 A display of genome compositional dynamics of SARS-CoV, MERS-CoV and their within-population CoVs** We use a 300-bp sliding window with a 21-bp step to show dynamic changes of genome G+C, purine, GC1 to GC3, and GC skew (G-C/G+C) contents. The complete genome sequences and data sources are listed in Table S1.

**Supplementary Figure S2 A**. G+C and purine content plot to show how these contents distribute among human CoVs. Note that all older human CoVs are drifted toward lower G+C and purine contents, and this phenomenon indicates lower selection pressure or insensitivity on composition changes. Full names of the human CoVs are listed in the legend of Figure 2. **B**. A genetic code table to show how nucleotide permutations are related to codons. The table is divided into two halves (colored and uncolored backgrounds) and cp1 and cp2 relative to their permutations sensitivity and changes are indicated with half parentheses with color-coding: C-to-U|U-to-C and A-to-G|G-to-A, red; G-to-U|U-to-G and A-to-C and C-to-U, blue; and A-to-U|U-to-A and G-to-C|C-to-G, green. Note that cp1 and cp2 are sensitive to column and row codon swaps, respectively. Cp3 is in a unique position where only half of the codons are sensitive to its changes, and the other half is so organized that some codons are more permissive than others.

**Supplementary Figure S3 Mutation spectra of SARS-CoV-2 in a context of codon positions A**. MERS-CoV and SARS-CoV mutation spectra are composed of 12 permutations and they are divided by codon positions. C-to-U (CU) permutations are always as dominant as what SARS-CoV-2 shows. The mutation counts are partitioned into synonymous and non-synonymous mutations. **B**. C-to-U permutations at cp3 diminish among non-synonymous mutations and this phenomenon indicates that most protein composition relevant variations are cp1 and cp2 variations. Note that all older human CoVs are drifted toward lower G+C and purine contents, and this phenomenon indicates lower selection or insensitivity on composition changes. Full names of the human CoVs are listed in the legend of Figure 2. Mutation counts are calculated from non-synonymous mutations. **C**. Displays of permutation-to-codon changes among non-synonymous mutations. The permutations showed here contain U-to-C, G-to-A, G-to-U, C-to-A, U-to-G, A-to-C, A-to-U, G-to-C, U-to-A and C-to-G.

**Supplementary Figure S4 High-frequency within-clade mutations** Signature site information and frequency table of star mutations in each clade, with a neighbor-joining tree based on the frequency data in the table.

**Supplementary Figure S5** This table illustrates how 12 permutations are related to G+C and purine contents and the subtle RTC specificity of CoVs. Mutations occur when the positive-sense RNA genome (R1, mutation happens when the negative-sense genome is synthesized) or its negative-sense subgenomes are synthesized (R2). The G+C content insensitive permutations occur after two syntheses (R12). CU and A-to-G (AG) are preferred when RTC is in a status that encourages a large-to-small substrate exchange in a higher ratio than the opposite, small-to-large substrate exchange. We term this preferred exchange as “tight” status. This status is also divided into R-tight (R, purine; to indicate that the mechanism is an A-by-G replacement) and Y-tight (Y, pyrimidine; to indicate that the mechanism is a C-by-U replacement). When an exchange of substrate happens from small-to-large, it is referred as a “loose” status that is also divided into two, R-loose and Y-loose. Note that some permutations are not sensitive to G+C and purine contents but others are sensitive. Arrow-headed dashed lines connect R1 permutation to R12 permutations, and note that cross-column relationship is rather striking, which re-routes some structural principles, which navigates mutation forces on one hand and leaves room for selection to work on, on the other hand.

**Supplementary Figure S6 Within-clade permutation variations are excellent indicators of functional mutations** (**A**) – (**D**) show within-clade permutation changes of C06, C08, C09 and C01.

**Supplementary Table S1 The mutation counts in each clade and cluster**

**Supplementary Table S2 The proportion of permutations in each clade and clusters based on different genomic regions**

**Supplementary Table S3 Selected CoV genome sequences used for this study Supplementary Table S4 The SARS-Cov-2 mutation table on coding region (based on data on June 12th 2020)**

**Supplementary Table S5 The proportion of permutations in each clade and their subclades**

## References

[1] Wu F, Zhao S, Yu B, Chen Y-M, Wang W, Song Z-G, et al. A new coronavirus associated with human respiratory disease in China. Nature 2020;579:265–9.

[2] World Health Organization. Coronavirus disease (COVID-2019) situation report - 144. https://www.who.int/emergencies/diseases/novel-coronavirus-2019/situation-reports (Jun 12 2020, date last accessed).

[3] He X, Lau EHY, Wu P, Deng X, Wang J, Hao X, et al. Temporal dynamics in viral shedding and transmissibility of COVID-19. Nat Med 2020;26:672–5.

[4] Zou L, Ruan F, Huang M, Liang L, Huang H, Hong Z, et al. SARS-CoV-2 Viral Load in Upper Respiratory Specimens of Infected Patients. N Engl J Med 2020;382:1177–9.

[5] Wu Z, McGoogan JM. Characteristics of and Important Lessons From the Coronavirus Disease 2019 (COVID-19) Outbreak in China: Summary of a Report of 72314 Cases From the Chinese Center for Disease Control and Prevention. JAMA 2020.

[6] Coronaviridae Study Group of the International Committee on Taxonomy of Viruses. The species Severe acute respiratory syndrome-related coronavirus: classifying 2019-nCoV and naming it SARS-CoV-2. Nat Microbiol 2020;5:536–44.

[7] Zhu N, Zhang D, Wang W, Li X, Yang B, Song J, et al. A Novel Coronavirus from Patients with Pneumonia in China, 2019. N Engl J Med 2020;382:727–33.

[8] Cui J, Li F, Shi Z-L. Origin and evolution of pathogenic coronaviruses. Nat Rev Microbiol 2019;17:181–92.

[9] de Wit E, van Doremalen N, Falzarano D, Munster VJ. SARS and MERS: recent insights into emerging coronaviruses. Nat Rev Microbiol 2016;14:523–34.

[10] Fung TS, Liu DX. Human Coronavirus: Host-Pathogen Interaction. Annu Rev Microbiol 2019;73:529–57.

[11] Yu J. From Mutation Signature to Molecular Mechanism in the RNA World: A Case of SARS-CoV-2. Genomics Proteomics Bioinformatics 2020;in press.

[12] Guo Y-R, Cao Q-D, Hong Z-S, Tan Y-Y, Chen S-D, Jin H-J, et al. The origin, transmission and clinical therapies on coronavirus disease 2019 (COVID-19) outbreak - an update on the status. Mil Med Res 2020;7:11.

[13] Smith EC, Blanc H, Surdel MC, Vignuzzi M, Denison MR. Coronaviruses lacking exoribonuclease activity are susceptible to lethal mutagenesis: evidence for proofreading and potential therapeutics. PLoS Pathog 2013;9:e1003565.

[14] Simmonds P. Rampant C-->U Hypermutation in the Genomes of SARS-CoV-2 and Other Coronaviruses: Causes and Consequences for Their Short- and Long-Term Evolutionary Trajectories. mSphere 2020;5.

[15] Di Giorgio S, Martignano F, Torcia MG, Mattiuz G, Conticello SG. Evidence for host-dependent RNA editing in the transcriptome of SARS-CoV-2. Sci Adv 2020;6:eabb5813.

[16] Xiao J, Yu J. A scenario on the stepwise evolution of the genetic code. Genomics Proteomics Bioinformatics 2007;5:143–51.

[17] Yu J. A content-centric organization of the genetic code. Genomics Proteomics Bioinformatics 2007;5:1–6.

[18] Zhang Z, Yu J. On the organizational dynamics of the genetic code. Genomics Proteomics Bioinformatics 2011;9:21–9.

[19] Zhang Z, Yu J. The pendulum model for genome compositional dynamics: from the four nucleotides to the twenty amino acids. Genomics Proteomics Bioinformatics 2012;10:175–80.

[20] DeWitte-Orr SJ, Collins SE, Bauer CMT, Bowdish DM, Mossman KL. An accessory to the ‘Trinity’: SR-As are essential pathogen sensors of extracellular dsRNA, mediating entry and leading to subsequent type I IFN responses. PLoS Pathog 2010;6:e1000829.

[21] Totura AL, Baric RS. SARS coronavirus pathogenesis: host innate immune responses and viral antagonism of interferon. Curr Opin Virol 2012;2:264–75.

[22] Crick FH. Codon--anticodon pairing: the wobble hypothesis. J Mol Biol 1966;19:548–55.

[23] Cui P, Lin Q, Ding F, Hu S, Yu J. The transcript-centric mutations in human genomes. Genomics Proteomics Bioinformatics 2012;10:11–22.

[24] Cui P, Ding F, Lin Q, Zhang L, Li A, Zhang Z, et al. Distinct contributions of replication and transcription to mutation rate variation of human genomes. Genomics Proteomics Bioinformatics 2012;10:4–10.

[25] Wong GK-S, Wang J, Tao L, Tan J, Zhang J, Passey DA, et al. Compositional gradients in Gramineae genes. Genome Res 2002;12:851–6.

[26] Lobry JR. Asymmetric substitution patterns in the two DNA strands of bacteria. Mol Biol Evol 1996;13:660–5.

[27] Wu Z, Lu L, Du J, Yang L, Ren X, Liu B, et al. Comparative analysis of rodent and small mammal viromes to better understand the wildlife origin of emerging infectious diseases. Microbiome 2018;6:178.

[28] Pan Y, Zhang D, Yang P, Poon LLM, Wang Q. Viral load of SARS-CoV-2 in clinical samples. Lancet Infect Dis 2020;20:411–2.

[29] Wang W, Xu Y, Gao R, Lu R, Han K, Wu G, et al. Detection of SARS-CoV-2 in Different Types of Clinical Specimens. JAMA 2020.

[30] Rambaut A, Holmes EC, O’Toole A, Hill V, McCrone JT, Ruis C, et al. A dynamic nomenclature proposal for SARS-CoV-2 lineages to assist genomic epidemiology. Nat Microbiol 2020.

[31] Tang X, Wu C, Li X, Song Y, Yao X, Wu X, et al. On the origin and continuing evolution of SARS-CoV-2. National Science Review 2020.

[32] Elbe S, Buckland-Merrett G. Data, disease and diplomacy: GISAID’s innovative contribution to global health. Glob Chall 2017;1:33–46.

[33] Zhou H, Chen X, Hu T, Li J, Song H, Liu Y, et al. A Novel Bat Coronavirus Closely Related to SARS-CoV-2 Contains Natural Insertions at the S1/S2 Cleavage Site of the Spike Protein. Curr Biol 2020;30:2196–203 e3.

[34] Zhou P, Yang XL, Wang XG, Hu B, Zhang L, Zhang W, et al. A pneumonia outbreak associated with a new coronavirus of probable bat origin. Nature 2020;579:270–3.

[35] Becerra-Flores M, Cardozo T. SARS-CoV-2 viral spike G614 mutation exhibits higher case fatality rate. Int J Clin Pract 2020:e13525.

[36] Daniloski Z, Guo X, Sanjana NE. The D614G mutation in SARS-CoV-2 Spike increases transduction of multiple human cell types. bioRxiv 2020:2020.06.14.151357.

[37] Korber B, Fischer W, Gnanakaran S, Yoon H, Theiler J, Abfalterer W, et al. Spike mutation pipeline reveals the emergence of a more transmissible form of SARS-CoV-2. bioRxiv 2020:2020.04.29.069054.

[38] Zhang L, Jackson CB, Mou H, Ojha A, Rangarajan ES, Izard T, et al. The D614G mutation in the SARS-CoV-2 spike protein reduces S1 shedding and increases infectivity. bioRxiv 2020:2020.06.12.148726.

[39] Wang M, Fu A, Hu B, Tong Y, Liu R, Gu J, et al. Nanopore target sequencing for accurate and comprehensive detection of SARS-CoV-2 and other respiratory viruses. medRxiv 2020:2020.03.04.20029538.

[40] Teymoori-Rad M, Samadizadeh S, Tabarraei A, Moradi A, Shahbaz MB, Tahamtan A. Ten challenging questions about SARS-CoV-2 and COVID-19. Expert Rev Respir Med 2020.

[41] Liu Q, Zhao S, Shi C-M, Song S-H, Zhu S, Su Y, et al. Population genetics of SARS-CoV-2: disentangling sampling bias and clustering infections. Genomics Proteomics Bioinformatics 2020;in press.

[42] Cotten M, Watson SJ, Kellam P, Al-Rabeeah AA, Makhdoom HQ, Assiri A, et al. Transmission and evolution of the Middle East respiratory syndrome coronavirus in Saudi Arabia: a descriptive genomic study. Lancet 2013;382:1993–2002.

[43] Gire SK, Goba A, Andersen KG, Sealfon RSG, Park DJ, Kanneh L, et al. Genomic surveillance elucidates Ebola virus origin and transmission during the 2014 outbreak. Science 2014;345:1369–72.

[44] Lemey P, Suchard M, Rambaut A. Reconstructing the initial global spread of a human influenza pandemic: A Bayesian spatial-temporal model for the global spread of H1N1pdm. PLoS Curr 2009;1:RRN1031.

[45] Smith GJD, Vijaykrishna D, Bahl J, Lycett SJ, Worobey M, Pybus OG, et al. Origins and evolutionary genomics of the 2009 swine-origin H1N1 influenza A epidemic. Nature 2009;459:1122–5.

[46] Yu W-B, Tang G-D, Zhang L, Corlett RT. Decoding the evolution and transmissions of the novel pneumonia coronavirus (SARS-CoV-2 / HCoV-19) using whole genomic data. Zool Res 2020;41:247–57.

[47] Sanders W, Fritch EJ, Madden EA, Graham RL, Vincent HA, Heise MT, et al. Comparative analysis of coronavirus genomic RNA structure reveals conservation in SARS-like coronaviruses. bioRxiv 2020:2020.06.15.153197.

[48] National Genomics Data Center M, Partners. Database Resources of the National Genomics Data Center in 2020. Nucleic Acids Res 2020;48:D24–D33.

[49] Wang B, Liu F, Zhang EC, Wo CL, Chen J, Qian PY, et al. The China National GeneBank horizontal line owned by all, completed by all and shared by all. Hereditas(Beijing) 2019;41:761–72.

[50] Sayers EW, Cavanaugh M, Clark K, Ostell J, Pruitt KD, Karsch-Mizrachi I. GenBank. Nucleic Acids Res 2020;48:D84–D6.

[51] Wu L, Sun Q, Desmeth P, Sugawara H, Xu Z, McCluskey K, et al. World data centre for microorganisms: an information infrastructure to explore and utilize preserved microbial strains worldwide. Nucleic Acids Res 2017;45:D611–D8.

[52] Edgar RC. MUSCLE: multiple sequence alignment with high accuracy and high throughput. Nucleic Acids Res 2004;32:1792–7.

[53] Kumar S, Stecher G, Li M, Knyaz C, Tamura K. MEGA X: Molecular Evolutionary Genetics Analysis across Computing Platforms. Mol Biol Evol 2018;35:1547–9.

[54] Marra MA, Jones SJ, Astell CR, Holt RA, Brooks-Wilson A, Butterfield YS, et al. The Genome sequence of the SARS-associated coronavirus. Science 2003;300:1399–404.

[55] McLaren W, Gil L, Hunt SE, Riat HS, Ritchie GRS, Thormann A, et al. The Ensembl Variant Effect Predictor. Genome Biol 2016;17:122.

[56] Price MN, Dehal PS, Arkin AP. FastTree 2--approximately maximum-likelihood trees for large alignments. PLoS One 2010;5:e9490.

[57] Letunic I, Bork P. Interactive Tree Of Life (iTOL) v4: recent updates and new developments. Nucleic Acids Res 2019;47:W256–W9.

[58] Schliep KP. phangorn: phylogenetic analysis in R. Bioinformatics 2011;27:592–3.

[59] Yu G. Using ggtree to Visualize Data on Tree-Like Structures. Curr Protoc Bioinformatics 2020;69:e96.

