## Supplementary material for "Compositional Variability and Mutation Spectra of Monophyletic SARS-CoV-2 Clades": FigureS4.pdf

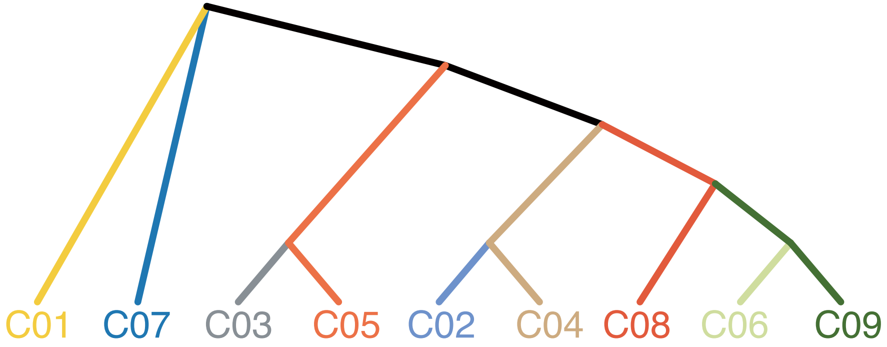


| Mutation | C01 | C07 | C03 | C05 | C02 | C04 | C08 | C06 | C09 | Protein | Mut Type |
|---|---|---|---|---|---|---|---|---|---|---|---|
| C241U | 0.1392 | 0 | 0.006 | 0.0342 | 0.0021 | 0 | 1 | 1 | 1 | 5'UTR | S |
| C1059U | 0.0466 | 0 | 0.0049 | 0 | 0.001 | 0 | 0.0006 | 0.0041 | 1 | nsp2 | NS |
| AAUG1604A | 0 | 1 | 0 | 0 | 0 | 0 | 0 | 0 | 0 | nsp2 | NS |
| C2416U | 0.0044 | 0 | 0.0005 | 0 | 0 | 0 | 0 | 0.079 | 0 | nsp2 | S |
| A2480G | 0.0036 | 0 | 0.0005 | 0 | 0 | 0 | 0.0002 | 0 | 0 | nsp2 | NS |
| C2558U | 0.0029 | 0 | 0.0005 | 0.0068 | 0 | 0 | 0.0002 | 0 | 0 | nsp2 | NS |
| C3037U | 0.1341 | 0.0024 | 0.0016 | 0 | 0.0052 | 0 | 1 | 1 | 1 | nsp3 | S |
| C8782U | 0.019 | 0.0024 | 0.0016 | 0 | 1 | 1 | 0.0002 | 0.0008 | 0 | nsp4 | S |
| G11083U | 0 | 0.0243 | 1 | 0 | 0.0186 | 0.014 | 0.0162 | 0.0105 | 0.0221 | nsp6 | NS |
| C13730U | 0.0087 | 0 | 0.1951 | 0 | 0 | 0 | 0 | 0.0067 | 0.0002 | RdRp | NS |
| C14408U | 0.1567 | 0 | 0.0038 | 0.0068 | 0.0031 | 0 | 1 | 1 | 1 | RdRp | NS |
| C14805U | 0.0058 | 0 | 0.5913 | 0.4041 | 0.2223 | 0.0007 | 0.0058 | 0 | 0 | RdRp | S |
| C15324U | 0.016 | 0 | 0 | 0 | 0.0041 | 0 | 0.0002 | 0.0922 | 0.0002 | RdRp | S |
| U17247C | 0.0015 | 0 | 0.2594 | 0.1301 | 0 | 0 | 0 | 0.0002 | 0 | helicase | S |
| C17747U | 0.0087 | 0 | 0.0005 | 0 | 0.0062 | 1 | 0.0004 | 0.0002 | 0 | helicase | NS |
| A17858G | 0.0087 | 0 | 0.0005 | 0 | 0.03 | 1 | 0 | 0 | 0 | helicase | NS |
| C18060U | 0.0124 | 0 | 0.0005 | 0 | 0.0424 | 1 | 0.0004 | 0.0002 | 0.0004 | xonucleas | S |
| C18877U | 0.0233 | 0 | 0.0005 | 0 | 0.0021 | 0.0007 | 0.001 | 0.1215 | 0 | xonucleas | S |
| A20268G | 0.0087 | 0 | 0 | 0 | 0.0031 | 0 | 0.0015 | 0.1725 | 0 | ndoRNAs | S |
| A23403G | 0.1895 | 0.0024 | 0.0027 | 0.0342 | 0.0062 | 0 | 1 | 1 | 1 | S | NS |
| G25563U | 0.0714 | 0 | 0.0033 | 0 | 0.0021 | 0.0007 | 0.0004 | 0.2499 | 1 | ORF3a | NS |
| G26144U | 0 | 0 | 0.6163 | 1 | 0.001 | 0 | 0 | 0 | 0 | ORF3a | NS |
| C27964U | 0.0051 | 0 | 0.0087 | 0 | 0 | 0 | 0 | 0.0005 | 0.1178 | ORF8 | NS |
| U28144C | 0.0481 | 0 | 0 | 0.0068 | 1 | 1 | 0 | 0 | 0.0004 | ORF8 | NS |
| C28311U | 0 | 0 | 0.0022 | 0 | 0 | 0 | 0 | 0 | 0 | N | NS |
| C28854U | 0.027 | 0.129 | 0.0005 | 0 | 0 | 0 | 0 | 0.0538 | 0.0087 | N | NS |
| G28881A | 0.0583 | 0.0024 | 0.0016 | 0.0205 | 0 | 0 | 1 | 0.0005 | 0.0008 | N | NS |
| G28882A | 0.0561 | 0 | 0 | 0.0137 | 0 | 0 | 1 | 0.0005 | 0.0006 | N | S |
| G28883C | 0.0561 | 0 | 0 | 0.0137 | 0 | 0 | 1 | 0.0005 | 0.0006 | N | NS |
