## Supplementary figures and images for "Compositional Variability and Mutation Spectra of Monophyletic SARS-CoV-2 Clades"

### FigureS1.pdf

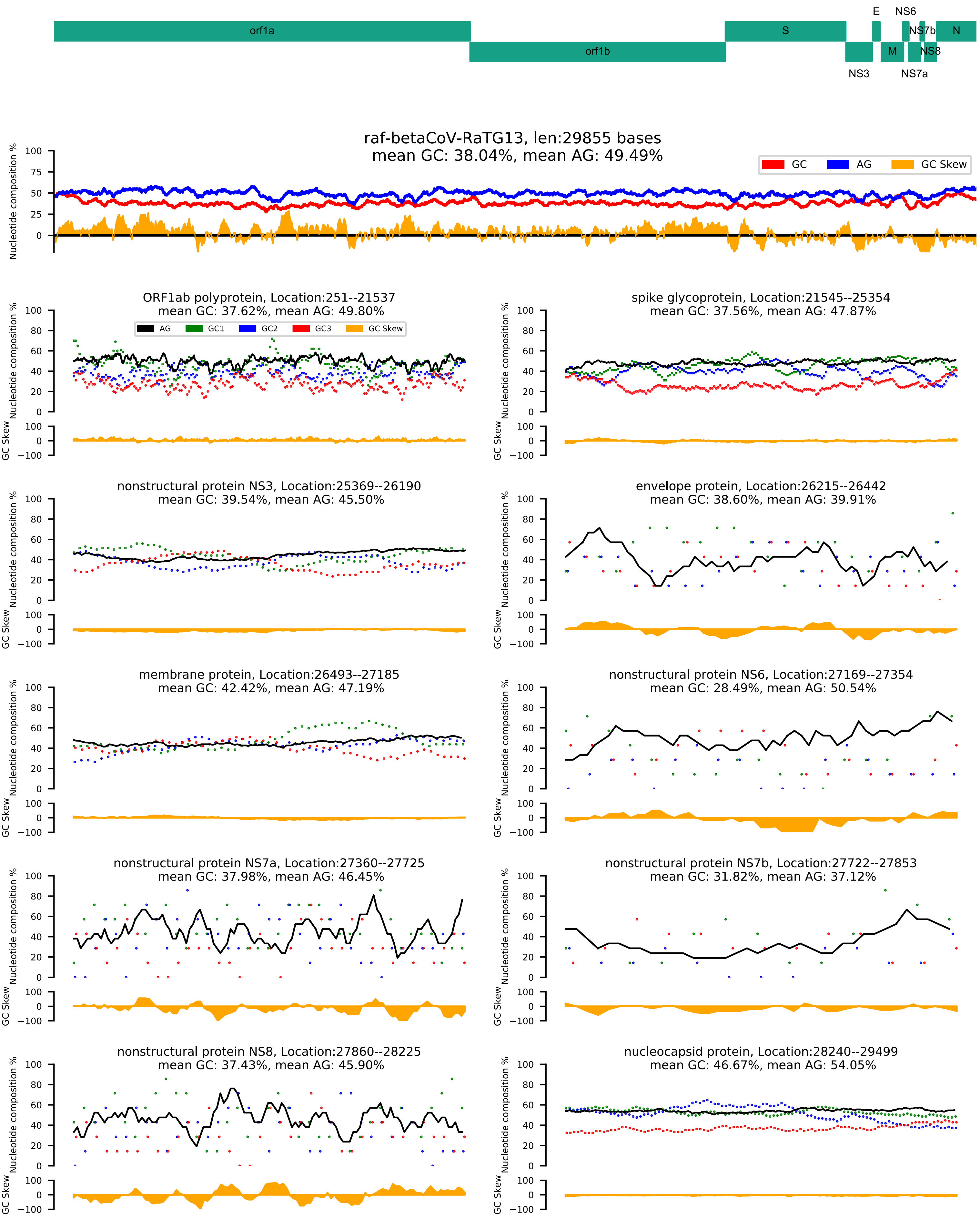

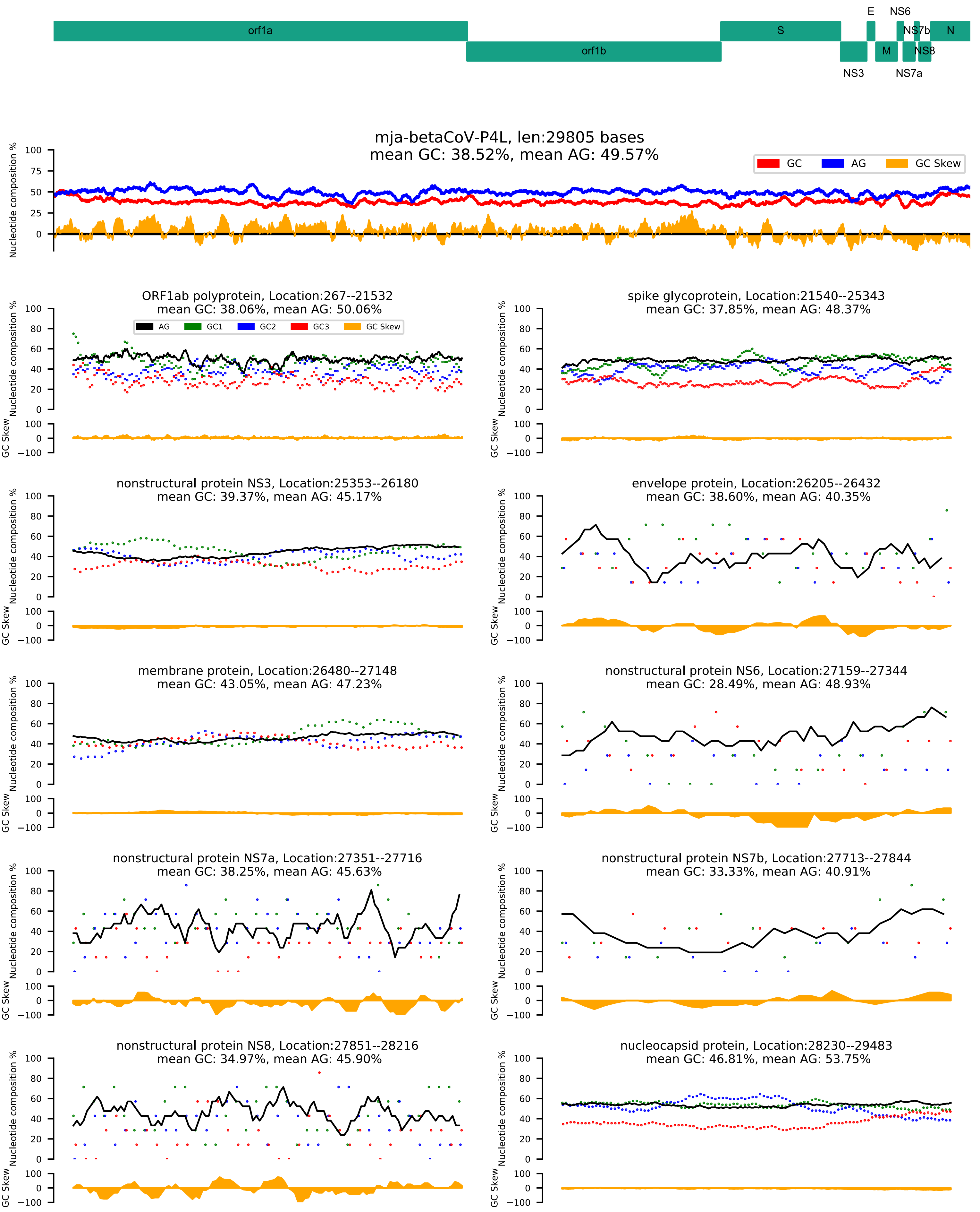

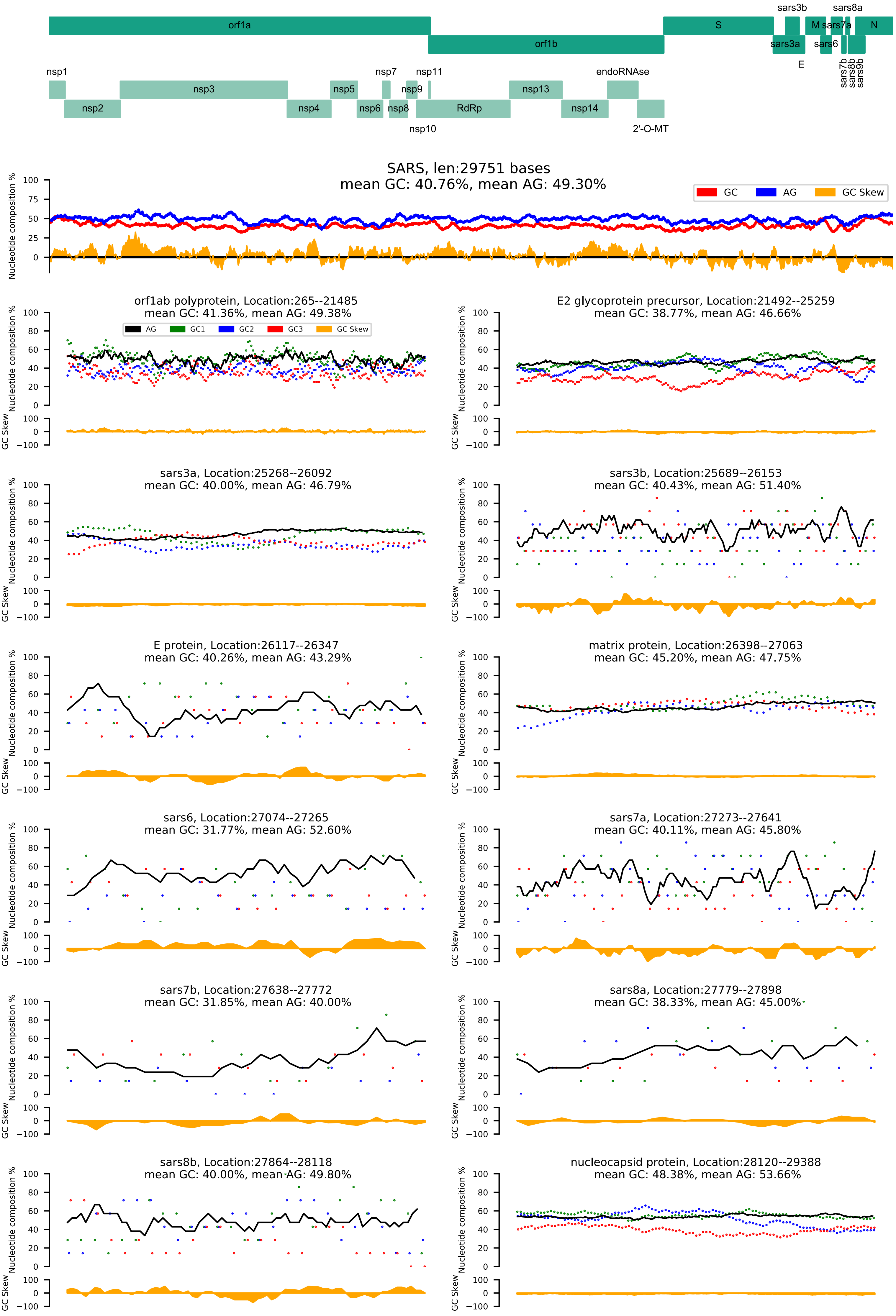

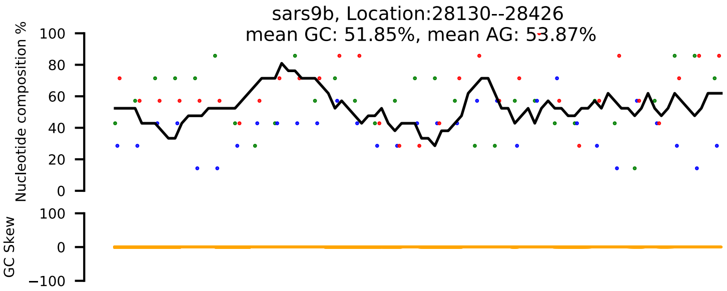

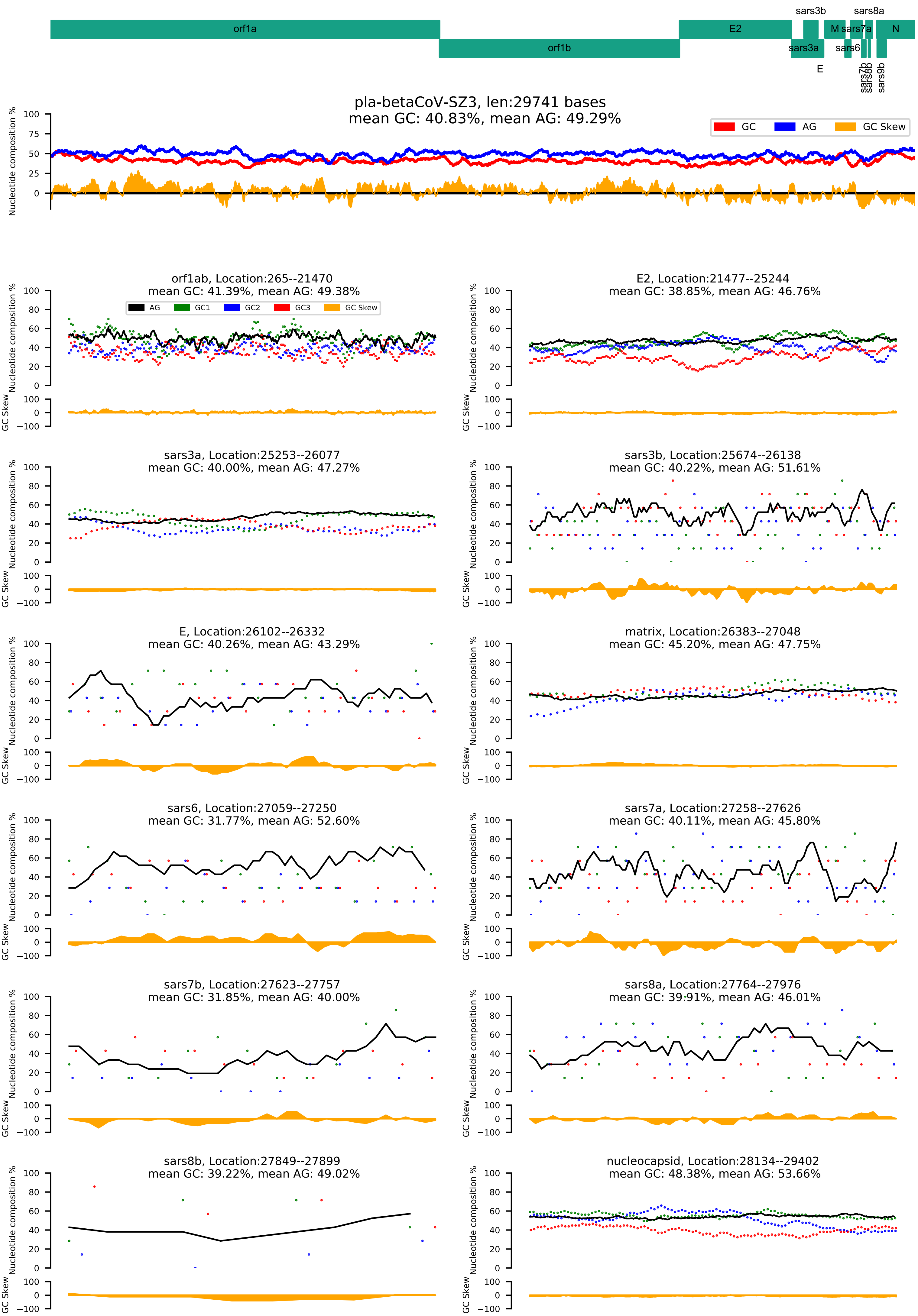

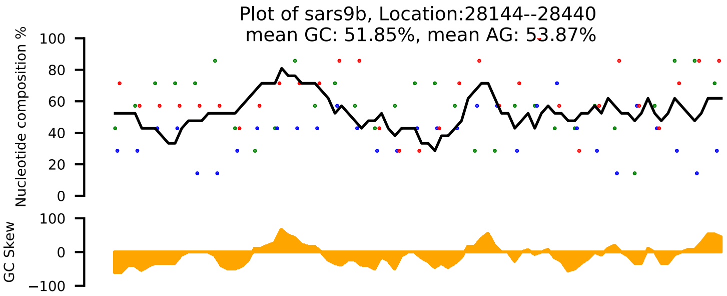

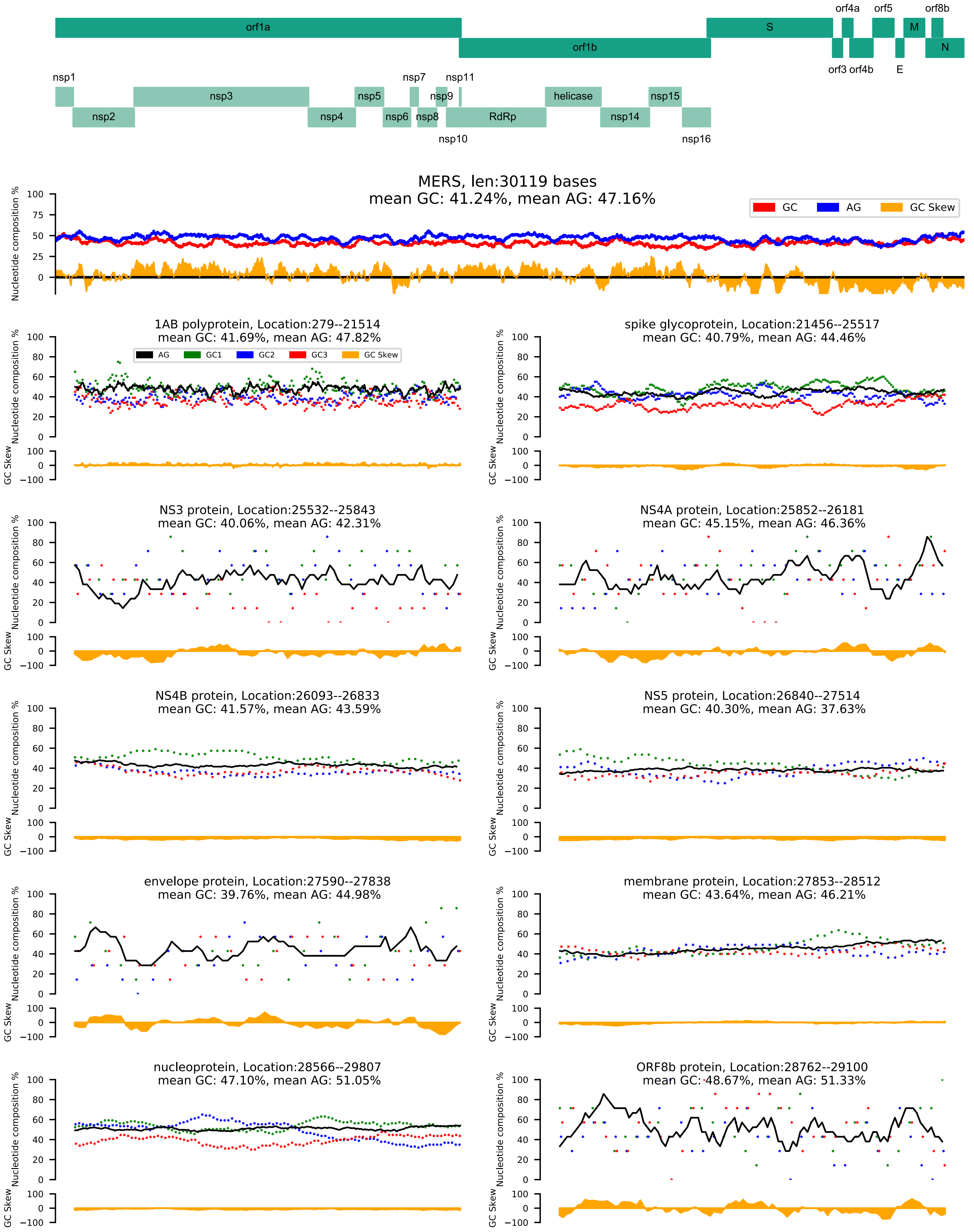

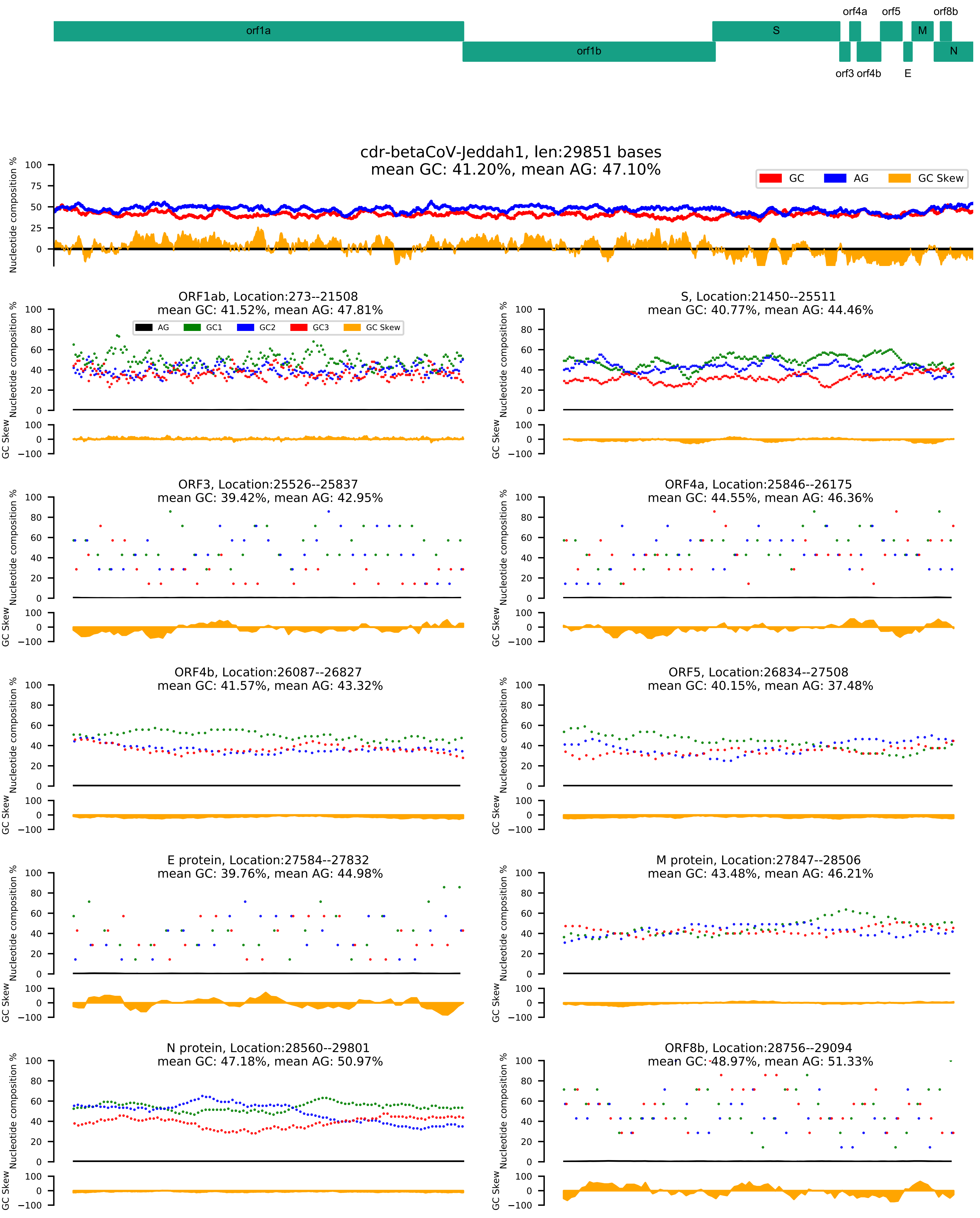

### FigureS2.pdf

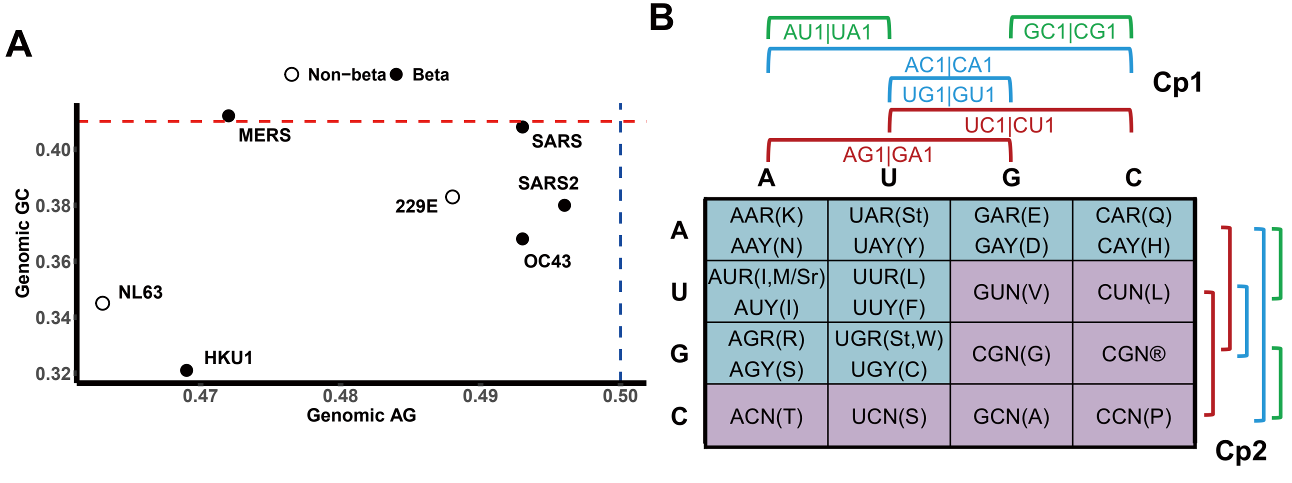

### FigureS3.pdf

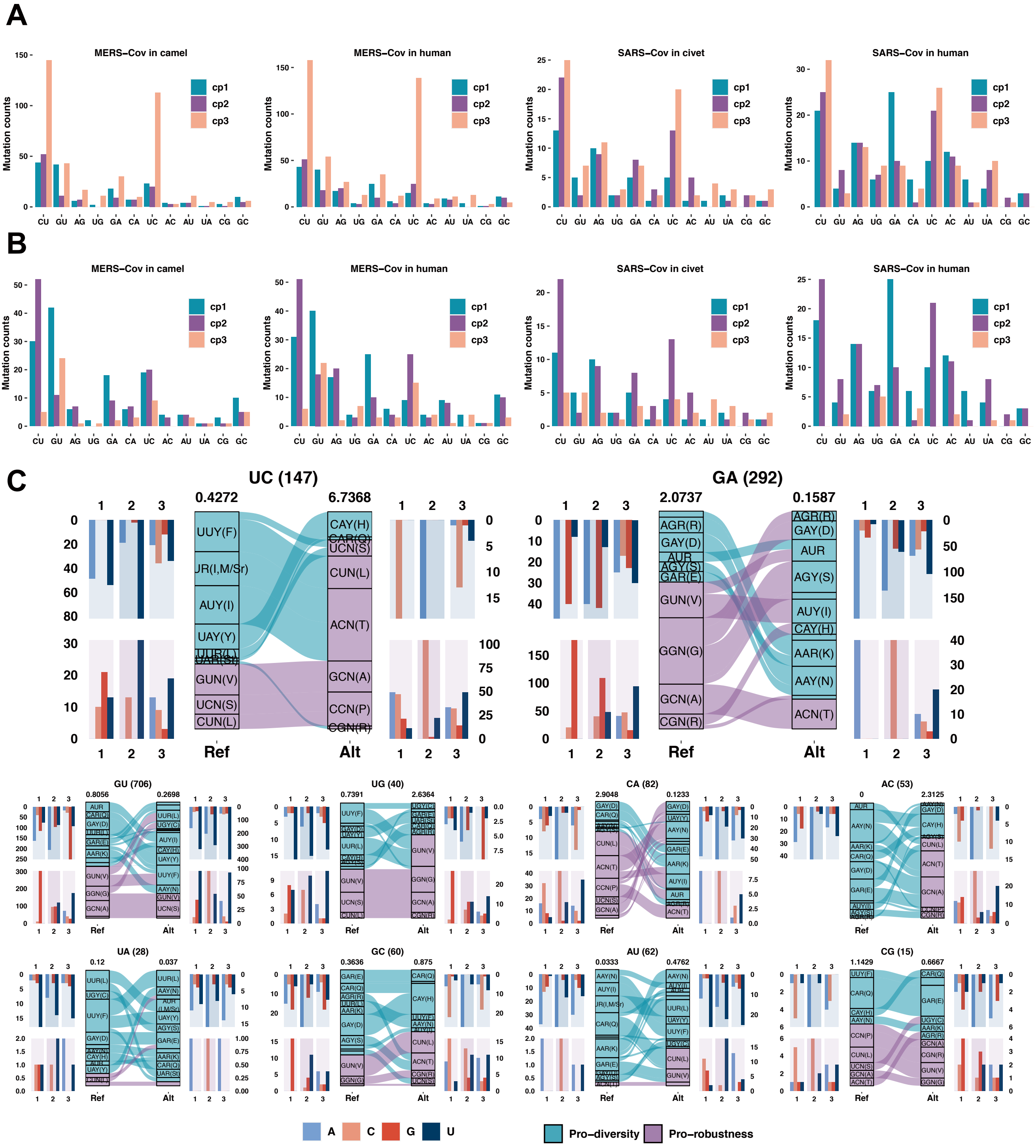

### FigureS5.pdf

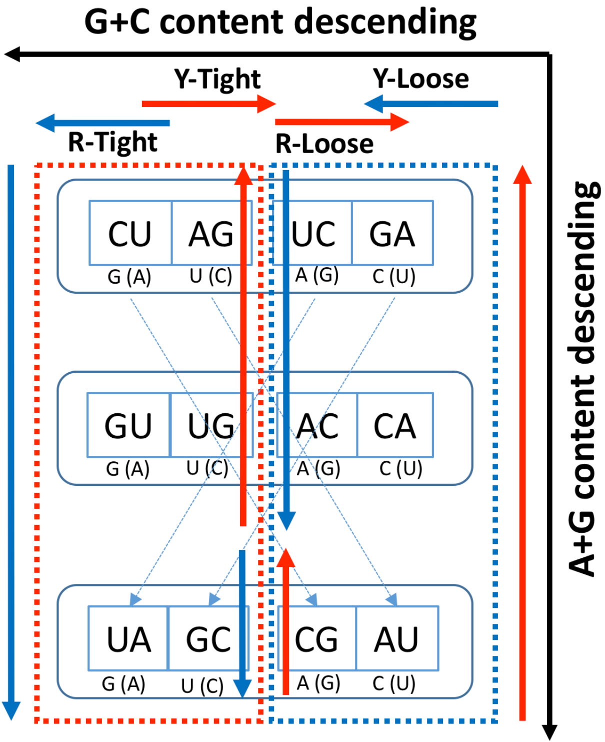

### FigureS6.pdf

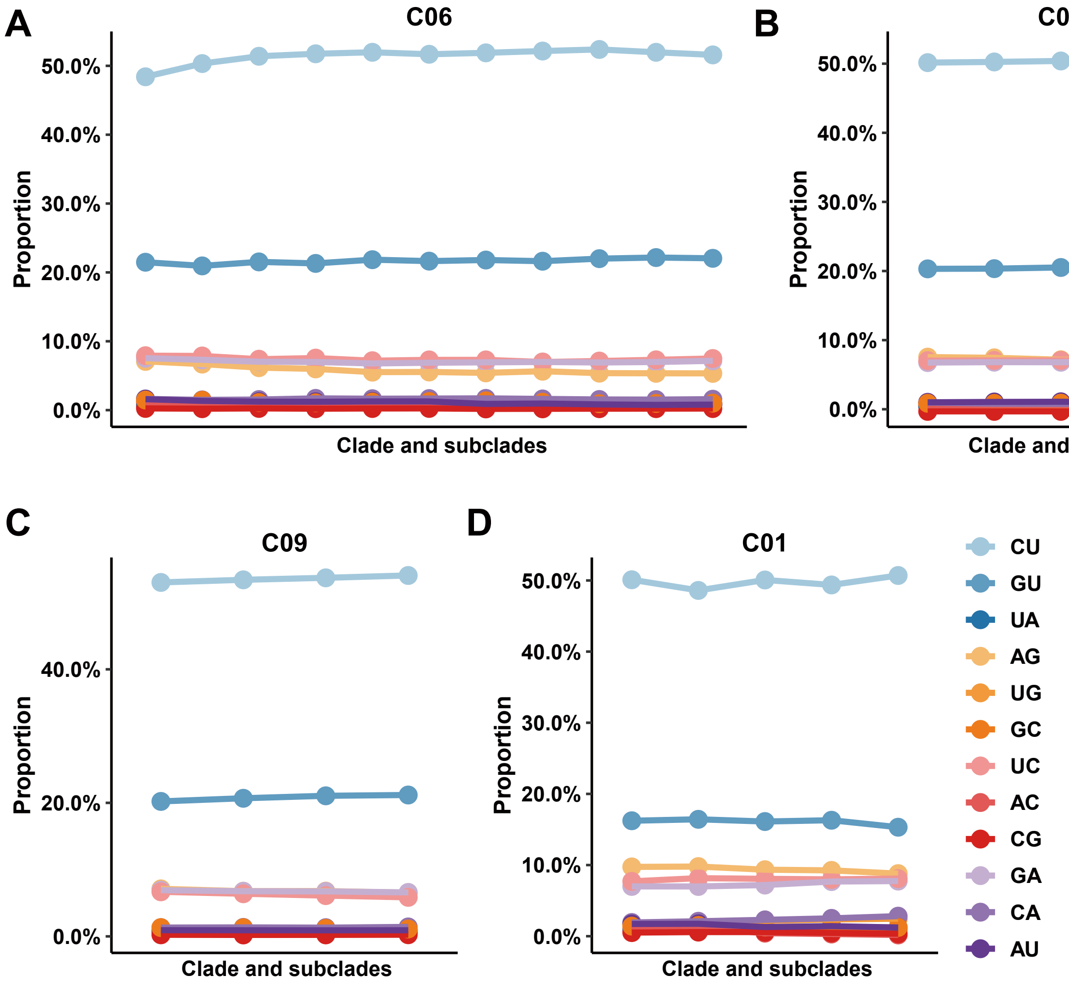
